# Systemic and local chronic inflammation and hormone disposition promote a tumor-permissive environment for breast cancer in older women

**DOI:** 10.1101/2024.10.18.616978

**Authors:** Neil Carleton, Sanghoon Lee, Ruxuan Li, Jian Zou, Daniel D Brown, Jagmohan Hooda, Alexander Chang, Rahul Kumar, Linda R Klei, Lora H Rigatti, Joseph Newsome, Dixcy Jaba Sheeba John Mary, Jennifer M Atkinson, Raymond E West, Thomas D Nolin, Patrick J Oberly, Ziyu Huang, Donald Poirier, Emilia J Diego, Peter C Lucas, George Tseng, Michael T Lotze, Priscilla F McAuliffe, Ioannis K Zervantonakis, Steffi Oesterreich, Adrian V Lee

**Affiliations:** Women’s Cancer Research Center, UPMC Hillman Cancer Center; Pittsburgh, PA, USA; Department of Bioengineering, University of Pittsburgh School of Medicine; Pittsburgh, PA, USA; Department of Statistics, School of Public Health, Chongqing Medical University; Chongqing, China; Department of Biostatistics, University of Pittsburgh School of Public Health; Pittsburgh, PA, USA; Institute of Precision Medicine, University of Pittsburgh Medical Center; Pittsburgh, PA, USA; Division of Laboratory Animal Resources, University of Pittsburgh School of Medicine; Pittsburgh, PA, USA; Department of Pathology, University of Pittsburgh School of Medicine; Pittsburgh, PA, USA; Small Molecule Biomarker Core, University of Pittsburgh School of Pharmacy; Pittsburgh, PA, USA; UPMC Hillman Cancer Center Biostatistics Facility; Pittsburgh, PA, USA; Laboratory of Medicinal Chemistry, CHU de Québec, Research Centre Université Laval, Quebec, Canada; Division of Breast Surgical Oncology, Department of Surgery, University of Pittsburgh School of Medicine; Pittsburgh, PA, USA; Department of Laboratory Medicine and Pathology, Mayo Clinic College of Medicine and Science; Rochester, MN, USA; Department of Immunology, University of Pittsburgh School of Medicine; Pittsburgh, PA, USA; Department of Pharmacology and Chemical Biology, University of Pittsburgh School of Medicine; Pittsburgh, PA, USA

**Keywords:** Aging, Estrogen Receptor Positive Breast Cancer, Older Patients with Breast Cancer, Chronic Inflammation, Estradiol, Estrogen Disposition

## Abstract

Estrogen receptor positive (ER+) breast cancer is the most common subtype of breast cancer and is an age-related disease. The peak incidence of diagnosis occurs around age 70, even though these post-menopausal patients have low circulating levels of estradiol (E2). Despite the hormone sensitivity of age-related tumors, we have a limited understanding of the interplay between systemic and local hormones, chronic inflammation, and immune changes that contribute to the growth and development of these tumors. Here, we show that aged F344 rats treated with the dimethylbenz(a)anthracene / medroxyprogestrone acetate (DMBA/MPA) carcinogen develop more tumors at faster rates than their younger counterparts, suggesting that the aged environment promotes tumor initiation and impacts growth. Single-nuclei RNA-seq (snRNA-seq) of the tumors showed broad local immune dysfunction that was associated with circulating chronic inflammation. Across a broad cohort of specimens from patients with ER+ breast cancer and age-matched donors of normal breast tissue, we observe that even with an estrone (E1)-predominant estrogen disposition in the systemic circulation, tumors in older patients increase *HSD17B7* expression to convert E1 to E2 in the tumor microenvironment (TME) and have local E2 levels similar to pre-menopausal patients. Concurrently, trackable increases in several chemokines, defined most notably by CCL2, promote a chronically inflamed but immune dysfunctional TME. This unique milieu in the aged TME, characterized by high local E2 and chemokine-enriched chronic inflammation, promotes both accumulation of tumor-associated macrophages (TAMs), which serve as signaling hubs, as well as polarization of TAMs towards a CD206+/PD-L1+, immunosuppressive phenotype. Pharmacologic targeting of estrogen signaling (either by HSD17B7 inhibition or with fulvestrant) and chemokine inflammation both decrease local E2 and prevent macrophage polarization. Overall, these findings suggest that chronic inflammation and hormonal disposition are critical contributors to the age-related nature of ER+ breast cancer development and growth and offer potential therapeutic insight to treat these patients.

**Translational Summary:** We uncover the unique underpinnings establishing how the systemic host environment contributes to the aged breast tumor microenvironment, characterized by high local estradiol and chronic inflammation with immune dysregulation, and show that targeting avenues of estrogen conversion and chronic inflammation work to restore anti-tumor immunity.

**Graphical Abstract:** 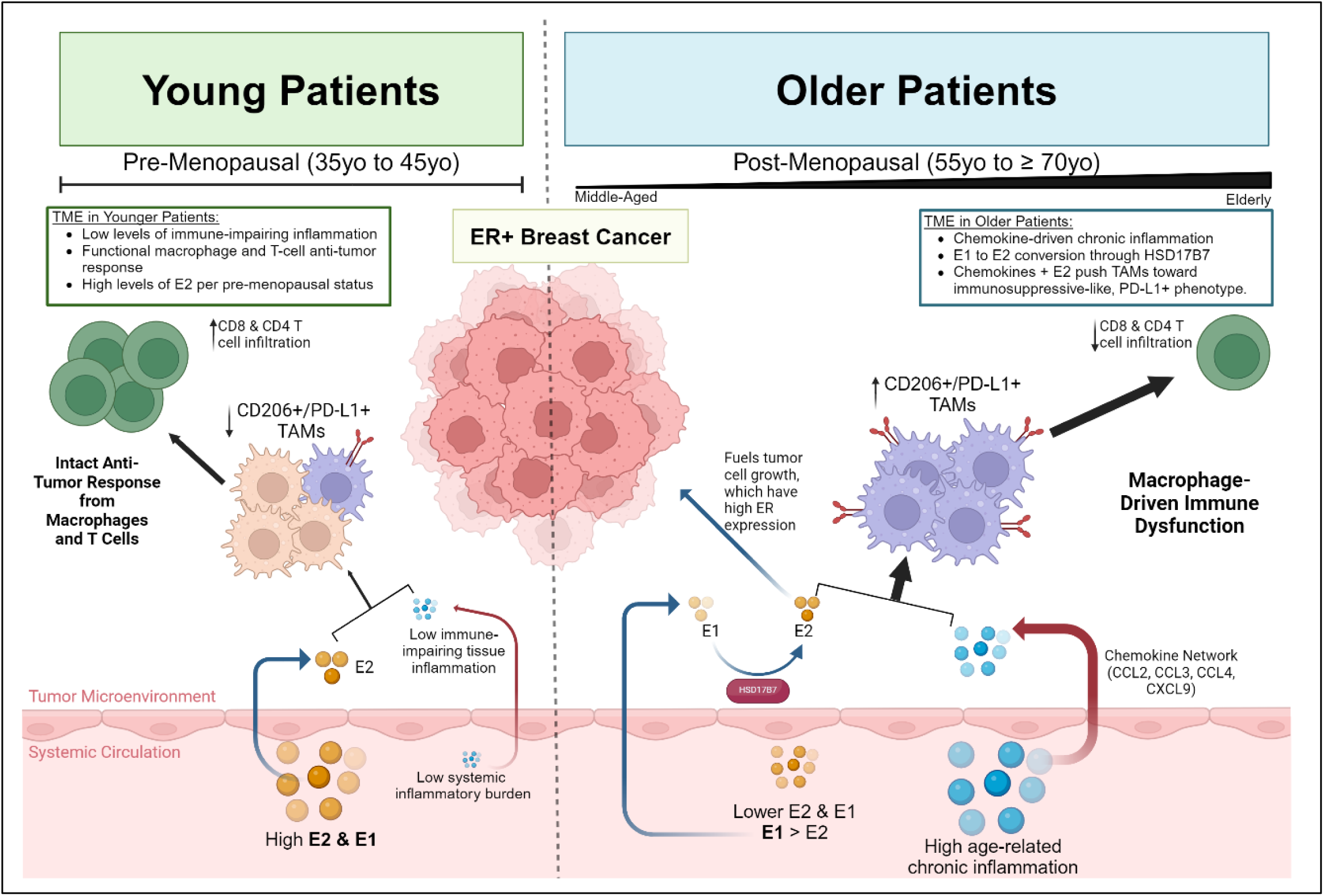

## INTRODUCTION

The biology of breast cancer is age-dependent (*1, 2*). Estrogen receptor positive (ER+) breast cancer, the most common subtype of breast cancer, is an age-related disease with the peak incidence of diagnosis occurring between the ages of 67 to 70 years old (*3*). This occurs despite low circulating estradiol (E2) levels for post-menopausal women. Further, the proportion of ER+ tumors continue to rise with advancing age: while ER+ breast cancer accounts for around 60%-70% of all tumors in women under 65, over 80%-90% of all breast tumors in those over 75 years old are considered ER+ (*4*).

While accumulating mutations are important contributors to the age-related nature of breast cancer (*5, 6*), the extent to which the environment – both systemic and within the breast itself – influences tumor development and growth are less well known. Tumors from older patients exhibit slightly higher tumor mutational burden compared to tumors in younger patients; however, many of the age-related changes in mutations reflect mutations that are typically associated with less aggressive, Luminal A-like tumors (*7, 8*). In fact, prior work minimizes these genomic changes in favor of the hypothesis that age-related breast tumors show some of the highest levels of transcriptomic alterations in comparison to other tumor types (*5, 9*). Tumors from older patients with ER+ disease are far less likely to exhibit gene expression changes indicative of faster growing tumors, such as growth factor receptor activation and p53 enrichment (*9, 10*). In a large study that included both breast cancer and normal breast tissues, the predominant age-correlated gene expression changes in breast cancers were attributable to changes in physiologic estrogen signaling, indicating a critical but unexplored role of how circulating and local factors, such as estrogen disposition and chronic inflammation, affect breast cancer development and growth in older women (*11*).

Along these lines, while cancer and aging are intricately linked from a cancer cell-intrinsic view, of increasing importance is the role of the tissue microenvironment and architecture, including stromal fibroblasts and adipocytes and the immune milieu, in supporting the development and growth of tumors (*12–14*). It is also becoming increasingly recognized that the systemic environment, most prominently defined by marked increase in chronic inflammation with age (*15, 16*), has a major impact on the development and growth of tumors in older patients (*17, 18*). In this bodywide ecological approach to understanding cancer, a person’s biological age, comorbidities, and comedications may alter systemic inflammation and hormone disposition, impacting immunosurveillance to new neoplastic lesions (*19*). The degree to which systemic factors shape the aged breast microenvironment is not well understood but is particularly important for how we craft clinical guidelines to better match the unique tumor biology in older patients.

Despite the prevalence of breast cancer in older patients, these patients remain under-represented not only in clinical trials but also in genomic and transcriptomic data sets (*20, 21*). This under-representation has led to a lack of preclinical and prospective data and, until recently, has precluded clinical guidelines and recommendations that reflected the behavior of these tumors in light of underlying biology and comorbidities (*22*). To this end, in this study, we aim to investigate how the systemic environment shapes the local aged breast microenvironment with a focus on changes in estrogen disposition and inflammation across young, middle-aged, and older patients. Using an aged rat model of carcinogen-induced ER+ breast cancer, we find that the aged rats develop more tumors and do so faster than their younger counterparts, in part driven by an inflamed but dysfunctional microenvironment. In a rich cohort of patient specimens, including analysis of matched blood and tissue from patients with ER+ breast cancer and donors of normal breast tumor, integrated analysis of TCGA, METABRIC, and SCAN-B, and analysis of publicly available scRNA-seq data, we find that an estrone (E1)- and chemokine-rich circulating phenotype converts to an E2-rich, chronically inflamed microenvironment with CD206+/PD-L1+ tumor associated macrophage-associated immune dysfunction. These data provide evidence supporting a tumor promoting role of the aged circulating and local breast environment that predisposes older patients to ER+ breast cancer development and growth.

## RESULTS

### Aged environment promotes tumors in older F344 rats in the DMBA/MPA carcinogen model

Fischer 344 (F344) inbred rats are one of the most commonly used models for aging, cancer, and toxicology studies. Despite this, relatively little has been reported on their estrogen levels, menstruation cycles, and age-related inflammation. We examined these age-related changes in estrogen disposition and inflammation in young (4 months old), middle-aged (12-13 months old), and older (20-22 months old) F344 rats. Four months of age corresponds to about the age of 20-30 years for humans while 21-23 months of age is the equivalent to about the age of 70 years for humans (**Fig 1A**). H&E-stained sections of ovary from young and older rats showed that the younger rats had active follicle formation while the older rats had no follicles present (**Fig. S1A**). Vaginal cytology for 14 consecutive days in six young and six older rats showed that young rats tended to progress through the estrus cycle every 5-6 days, while the older rats, with prominent and persistent pattern of cornified epithelial cells, were in a state of diestrus (**Fig. S1B**). This was concordant with the measured serum estradiol (E2) and estrone (E1) levels, which showed that older rats had significantly lower levels of E2 in a post-estropause state (**Fig. S1C**). Measurement of circulating inflammatory factors with an 18-plex assay revealed a chronic inflammatory state in the older rats characterized by increases in several chemokines TNFa, IL-1b, and IL-10 (**Fig. S1D**).

**Fig. 1:**
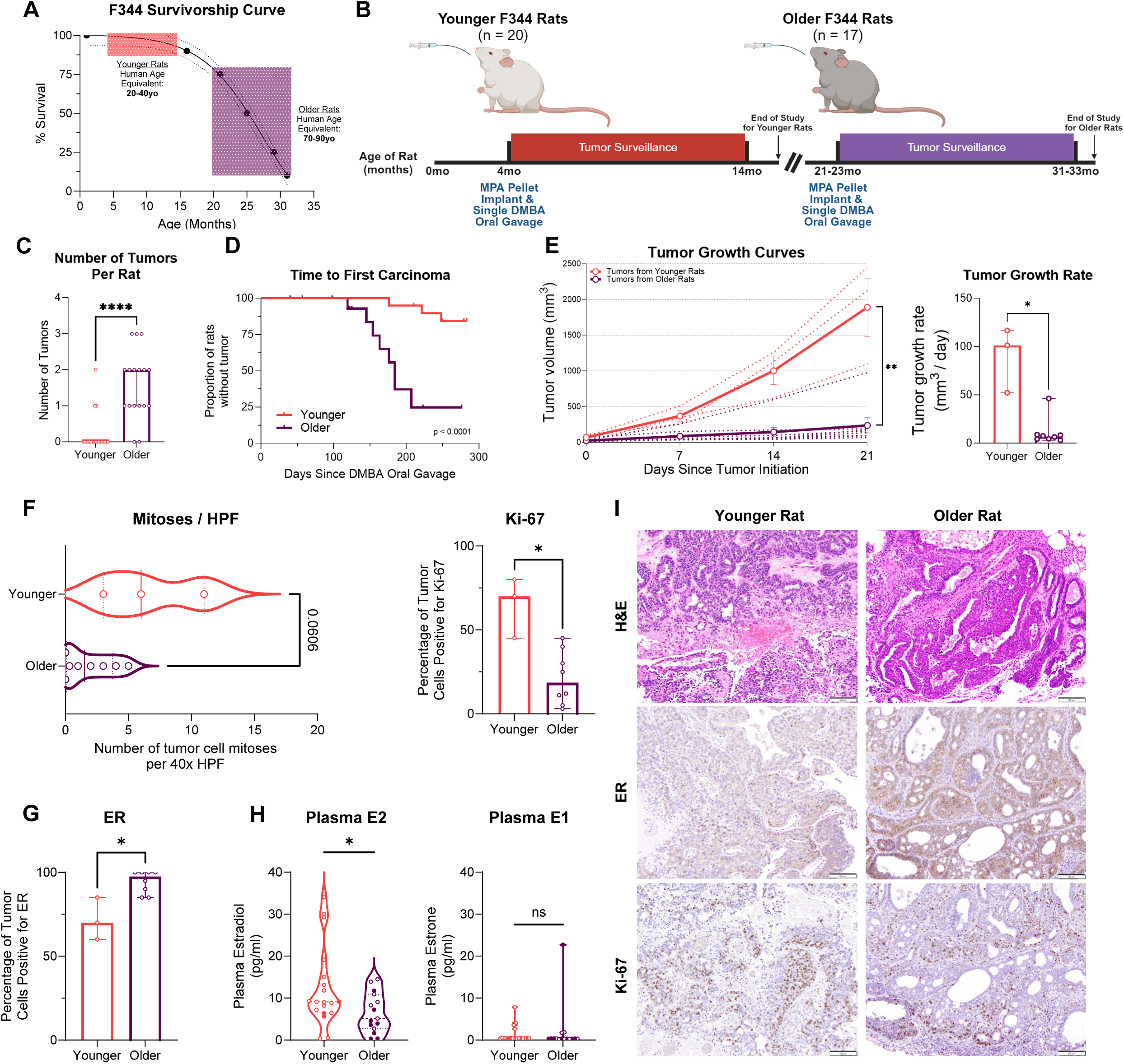
The aged environment promotes accelerated yet indolent tumorigenesis in a rat model of post-menopausal ER+ breast cancer. (A) Schematic of the experimental setup. In brief, younger F344 rats (n = 20, ∼ 4mo old) and older F344 rats (n = 17, ∼ 21-23mo old) were implanted with an MPA pellet and treated with a single dose of the DMBA carcinogen and were subsequently monitored for tumors over a 10mo time period. (B) Survivorship curve showing the human equivalent age for the younger and older rats used in the study. (C) Older rats harbored significantly more tumors than younger rats. This plot includes all tumors that formed, including both fibroepithelial lesions as well as mammary carcinomas. Data represent median and 95% CI; **** indicates p-value < 0.0001. (D) Time-to-event analysis showing the time to first carcinoma. Older rats had a significantly shorter time to first carcinoma compared to younger rats (p < 0.0001). Overall, there were three carcinomas in the younger rats and 8 carcinomas in the older rats. (E) Tumors in older rats generally grew slower when comparing both the tumor growth curves (all tumors compared at day 21; solid lines represent the average growth for each age group whereas the dotted lines show individual tumor trajectories) and the tumor growth rate (computed using the total tumor volume at the time of sacrifice). ** indicates p-value less than 0.01, * indicates p-value less than 0.05. (F) Shows mitoses per high power field (HPF) and the percentage of tumor cells staining positive for Ki-67; both data computed by a trained animal histopathologist using at least four different areas per tumor. Data show median and range for mitoses per HPF plot and median and 95% CI for Ki-67 plot. * indicates p-value less than 0.05. (G) Percent of tumor cells staining positive for estrogen receptor (ER). Data show median and 95% CI. * indicates p-value less than 0.05. (H) Plasma E2 and plasma E1 analyzed on isolated serum at the time of sacrifice for all younger (n = 20) and older rats (n = 17) in the study. Filled in dots indicate rats that had carcinomas at the time of sacrifice. Data show median with range. * indicates p-value less than 0.05. (I) Shows representative tumor images (H&Es, estrogen receptor [ER] staining, and Ki-67 staining) for younger and older rats.

This comprehensive characterization of estrogen levels, menstruation cycles, and age-related inflammation in F344 rats provided critical insights into their systemic environment, setting the stage for exploring how these factors influence tumor initiation. Based on this, to determine if and the degree to which the aged environment affects tumor development and growth, we utilized a carcinogen-induced model of ER+ breast cancer, which included a single oral gavage of the carcinogen 7,12-dimethylbenz[a]anthracene (DMBA) and an implanted slow-release pellet of medroxyprogesterone acetate (MPA). This model has been previously shown to produce estrogen sensitive tumors that rely on immune escape for tumor development and growth and generally have a prominent inflammatory reaction within the tumor microenvironment (*23–25*). Notably, many previous studies induced tumors in prepubescent rats (6-7 week of age) when they are most susceptible to carcinogen likely due to the developing cells in the mammary gland. In contrast, to better represent carcinogenesis in adult women, we used aged rats.

Younger rats (n = 20) were treated with the carcinogen at 4 months of age while older rats (n = 17) were treated at 21-23 months of age (**Fig. 1B**). Following treatment, older rats developed significantly more tumors, including both carcinomas and fibroepithelial lesions, than younger rats (p < 0.0001). After excluding the fibroepithelial lesions, the older rats had a significantly faster time to first carcinoma (p < 0.0001), a surprising finding given that rat mammary glands are thought to be less susceptible to carcinogen-induced carcinomas with advancing age (*24*) (**Figs. 1C and 1D**). Notably, tumors in older rats grew significantly slower than those in younger rats (p = 0.0015) and accordingly had fewer mitoses per 40x field (p = 0.06) and lower Ki-67+ tumor cells (p = 0.02) (**Figs. 1E and 1F**). Estrogen receptor (ER) expression was higher in the tumors from older rats (p = 0.02) despite lower circulating levels of estradiol (p = 0.04) (**Figs. 1G, 1H, and 1I**). These data suggest not only that the carcinogen-induced model recapitulates key features of age-related, post-menopausal breast cancer in humans, but also that the aged environment seems to promote increased tumorigenesis characterized by indolently-growing tumors in the older rats.

With these differences between tumors in younger and older rats, we utilized whole exome sequencing, bulk RNA-seq, and single-nuclei RNA-seq to further characterize age-related changes. Carcinogen-induced tumors showed a random mutational pattern across age groups with no clear age-correlated differences (**Fig. S2A**); when restricting to commonly mutated genes in human breast cancer, we again found little difference across age (**Fig. 2A**). COSMIC signature analysis revealed enrichment of SBS4 (signal of direct DNA mutagenesis) and SBS5 (clock-like signature in bladder cancer and other cancer types associated with tobacco smoking) across both age groups, likely reflecting the DMBA-induced nature of the tumors; SBS9 and SBS89, proposed to be markers of carcinogenesis in the first decade of life, were only present in tumors from young rats (**Fig. 2B**). Interestingly, tumor mutational burden (TMB) was decreased in tumors from older rats (p = 0.10; **Fig. 2C**). PAM50 subtyping of the tumors, which showed enrichment for mostly Luminal A-like intrinsic subtypes, did not vary by age (**Fig. 2D**). Single-nuclei RNA-seq of the tumors further confirmed minimal transcriptomic differences between in cancer epithelial cells in younger and older tumors (**Figs. 2F and S2B**). Given the lack of genomic and transcriptomic differences in tumors from younger and older rats, we further explored immune (single-nuclei RNA-seq) and inflammatory (circulating inflammation panel) differences. The most remarkable differences between tumors from younger and older rats were related to immune cell proportion and characterization as well as circulating inflammatory markers. Cells from the myeloid lineage generally increased with age (higher proportions in tumors from older rats; **Fig. 2E** with cell marker identification in **Figs. S2C and S2D**); yet these infiltrative macrophages in tumors from older rats exhibited lower levels of immune stimulating pathway enrichment (**Fig. 2F**). When we again analyzed an 18-plex circulating inflammatory panel from these rats, many of the markers were again higher in the older rats; interestingly, however, several chemokines that promote macrophage infiltration (CCL2, CCL3, GM-CSF, CXCL1, and CXCL10) were amongst the most significantly different between age groups (p < 0.05 across all markers; **Fig. 2G**). Overall, these data collectively suggests that the increase in tumor burden and tumor growth behavior with age is unlikely related to increased susceptibility to the carcinogen, differences in driver mutations with age that may confer phenotypic differences, or tumor mutational burden. Instead, we observe vast differences in the proportion and character of the infiltrating immune cells and chronic inflammatory phenotype in the older rats, suggesting a prominent role of macrophage-associated immune dysfunction in these tumors.

**Fig. 2:**
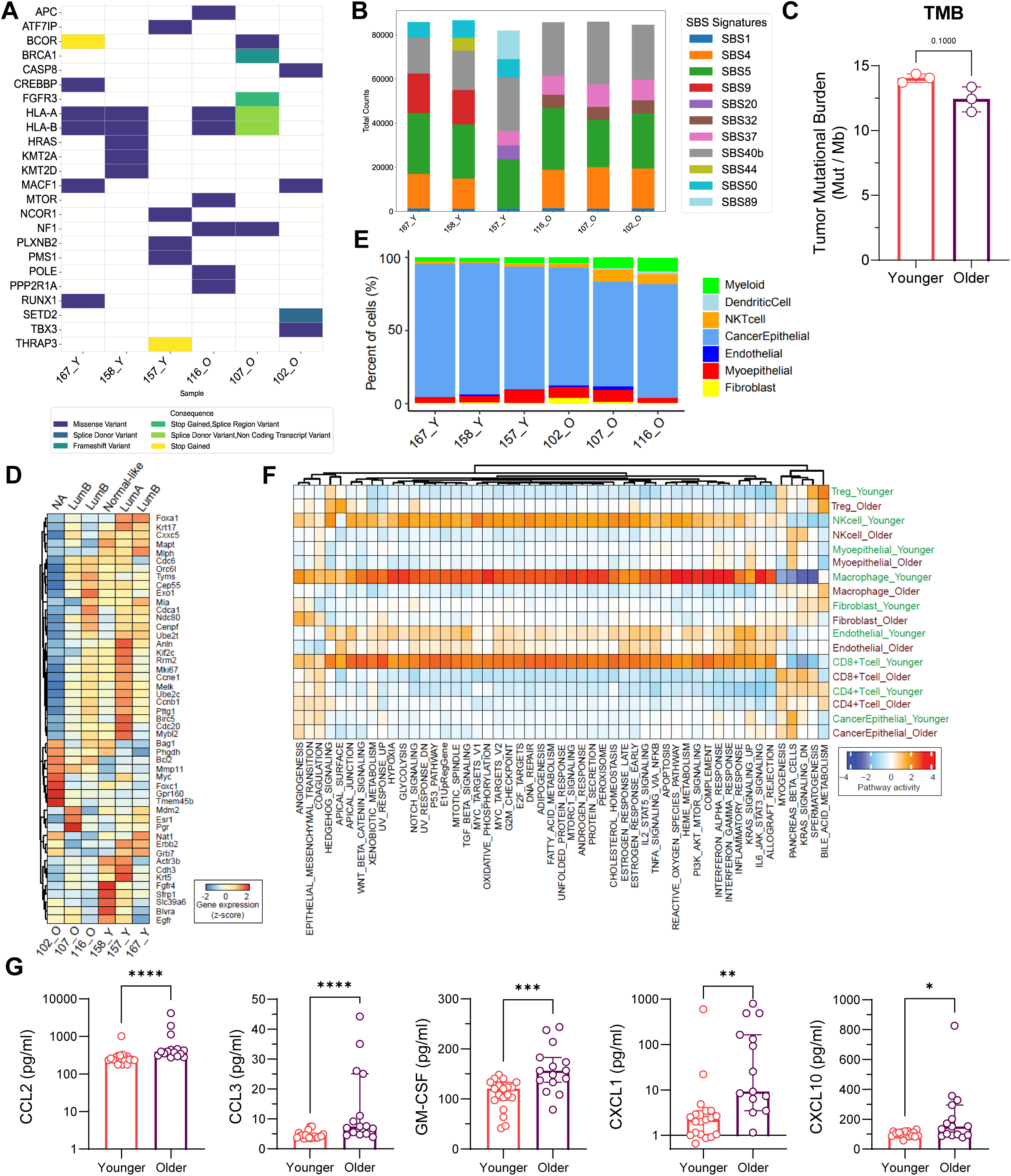
Accelerated tumorigenesis in older rats is unlikely related to intrinsic mechanisms but rather due to inflammation and immune differences. (A) Oncoplot for the three tumors from younger rats (labeled with tumor ID and “Y”) and three tumors from older rats (labeled with tumor ID and “O”) showing random mutations induced from the carcinogens without a pattern related to age of the host. (B) COSMIC signature analysis shows enrichment of SBS4 and SBS5 signatures across all tumors, related to the carcinogenic nature of the tumors. (C) Tumor mutational burden (TMB) was decreased in the tumors from older rats. (D) PAM50 analysis shows no distinct patterns by age; tumors aligned to Luminal A-like, Luminal B-like, and Normal-like. One tumor could not be assigned a PAM50 subtype. (E) Using snRNA-seq, we analyzed proportions of cells in the TME, showing that cells from the myeloid lineage were higher in the tumors from older rats compared to the tumors from younger rats. (F) Using the snRNA-seq data, we analyzed HALLMARK pathway expression across the immune cell subtypes in tumors from younger vs. older rats. (G) Cytokine analysis on the blood taken at the time of sacrifice (through cardiac puncture) shows significantly enriched levels of circulating chemokines in the older rats. Data show median and 95% CI. **** indicates p-value less than 0.0001, *** indicates p-value less than 0.001, ** indicates p-value less than 0.01, * indicates p-value less than 0.05.

### Aged circulating environment in humans is characterized by E1 predominance with chemokine-enriched chronic inflammation

We next constructed two cohorts of human specimens to investigate changes in both circulating and local factors with age: (1) matched plasma, tumor tissue, and tumor-adjacent tissue from n = 115 patients with ER+ breast cancer from the Pitt Biospecimen Core; and (2) matched plasma and noncancerous, normal breast tissue from n = 89 donors in the Komen Tissue Bank (KTB). For the patients with ER+ breast cancer, no patients received any neoadjuvant therapy and all blood and tissue specimens were collected at the time of surgery. We split the specimens into the following age groups: young, 35 to 45 years old; middle-aged, 55 to 69 years old; and older, ≥ 70 years old (**Fig. 3A**). While there are a variety of age cutoffs used to define “older” or “geriatric” in clinical practice, 70 was chosen as there are a number of clinical guidelines currently in place using this cutoff; in reality, aging is a heterogeneous process dependent on a multitude of factors and likely occurs at different rates in different people (*19, 26–29*). In general, there were no differences in tumor grade, stage, or histology amongst the three age groups (**Table S1**). In the cohort of donors from the Komen Tissue Bank, there were no differences in donor BMI, age at menarche, number of prior pregnancies, race, or ethnicity across age groups (**Table S2**). We note that a number of post-menopausal donors had previously used hormone replacement therapy, though none were using at the time of specimen donation.

**Fig. 3:**
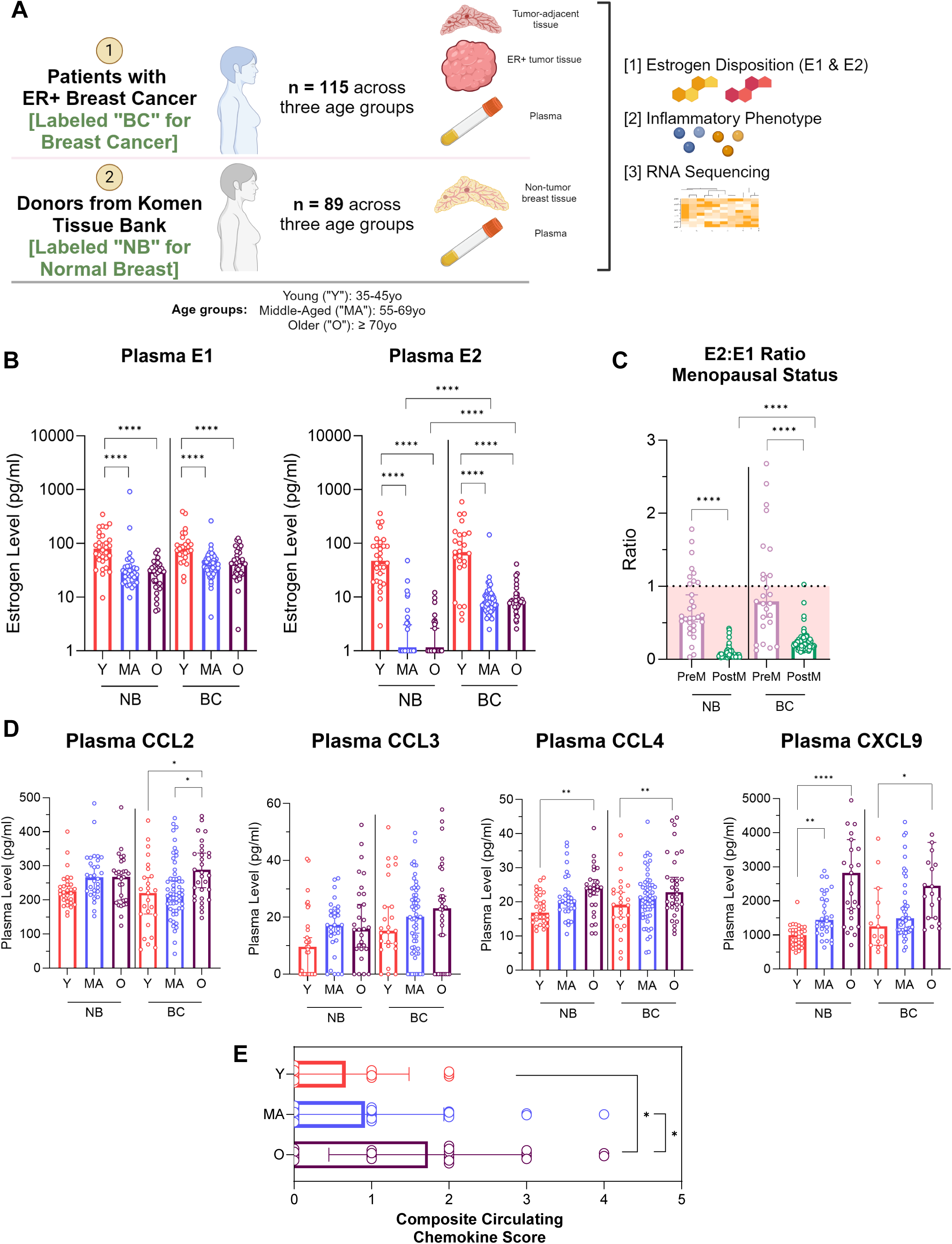
Systemic analysis of estrogen disposition and inflammatory phenotype shows an E1-predominant, chronically inflamed host environment. (A) Schematic of the key cohorts of human specimens utilized in the study: cohort of 115 patients with ER+ breast cancer with matched plasma, tumor tissue, and tumor-adjacent tissue; cohort of 90 donors from the Komen Tissue Bank at Indiana University with matched plasma and non-tumor, normal breast tissue. (B) Analysis of plasma E1 and E2 from both cohorts. NB indicates data derived from the donors in the Komen Tissue Bank; BC indicates data derived from patients with ER+ breast cancer. Age groups indicated by Y (young, 35-45yo), MA (middle-aged, 55-69yo), and O (older, ≥ 70yo). Data show median with 95% CI. **** indicates p-value less than 0.0001, data corrected for multiple comparisons. (C) E2:E1 ratio computed according to menopausal status. Data show median with 95% CI. **** indicates p-value less than 0.0001, data corrected for multiple comparisons. (D) Circulating levels of CCL2, CCL3, CCL4, and CXCL9. Data show median with 95% CI. **** indicates p-value less than 0.0001, ** indicates p-value less than 0.01, * indicates p-value less than 0.05. Data corrected for multiple comparisons. (E) Shows the computed composite circulating chemokine (scale 0-4, patient scored 1 point for each of the four chemokines in (D) that were above 75% of the measured values across age groups) score for each age group in those patients with ER+ breast cancer. Data indicate median with 95% CI. * indicates p-value less than 0.05. Data corrected for multiple comparisons.

Measurement of plasma E1 and E2 in donors without breast cancer and from patients with ER+ breast cancer showed that the loss of E2 in postmenopausal women causes E1 to become the predominant circulating estrogen (**Fig. 3B**), consistent with prior reports (*30*). There were no significant differences in E1 or E2 between the middle-aged and older groups of donors and patients. However, we noted elevated circulating E2 and the E2:E1 ratio in postmenopausal ER+ breast cancer patients compared to donors without breast cancer (**Figs. 3B and 3C**). From the same plasma samples, we measured a 22-plex panel of inflammatory factors.

Both donors without breast cancer and patients with ER+ breast cancer exhibited a marked increase in inflammation with increasing age (**Fig. S3**). Using adjusted linear regression, we identified a series of chemokines, CCL2, CCL3, CCL4, and CXCL9, that were most associated with increasing age across the plasma samples (**Fig. 3D**). These factors, and specifically, the combined burden of these markers, were highest in the plasma of older donors and patients (**Fig. 3E**). Collectively, these data suggests that while estrogen disposition is largely a product of menopausal status and does not change between middle-aged and older donors and patients, a chronic inflammatory phenotype, defined by a chemokine network, is enriched in both older donors and patients.

### Age-related *HSD17B7* expression contributes to E2 enrichment in the local breast TME in post-menopausal patients

We next investigated estrogen disposition in the normal breast microenvironment from KTB donors as well as in the TME from patients with breast cancer and their adjacent tissues. In the normal breast, E2 in the breast of older donors was significantly lower than E1 (p = 0.0033) (**Fig. 4A**), mirroring the difference seen in plasma. In stark contrast, despite the vast plasma differences in E1 and E2 with menopause, E2 was the predominant estrogen in the breast for patients with ER+ disease with marked increases over E1 across all age groups. In the tumor tissue itself, E2 was similar across all groups (p = 0.99) (**Fig. 4A**). Of note, we did observe differences in BMI across the cohorts (**Fig. S4A**): while BMI was expectedly associated with plasma E2 in post-menopausal women (**Fig. S4B**), it was also weakly associated with tissue E2 as well (**Fig. S4C**). There were no differences in plasma or tumor estrogen levels in patients with IDC versus those with ILC (**Figs. S5A and S5B**).

**Fig. 4:**
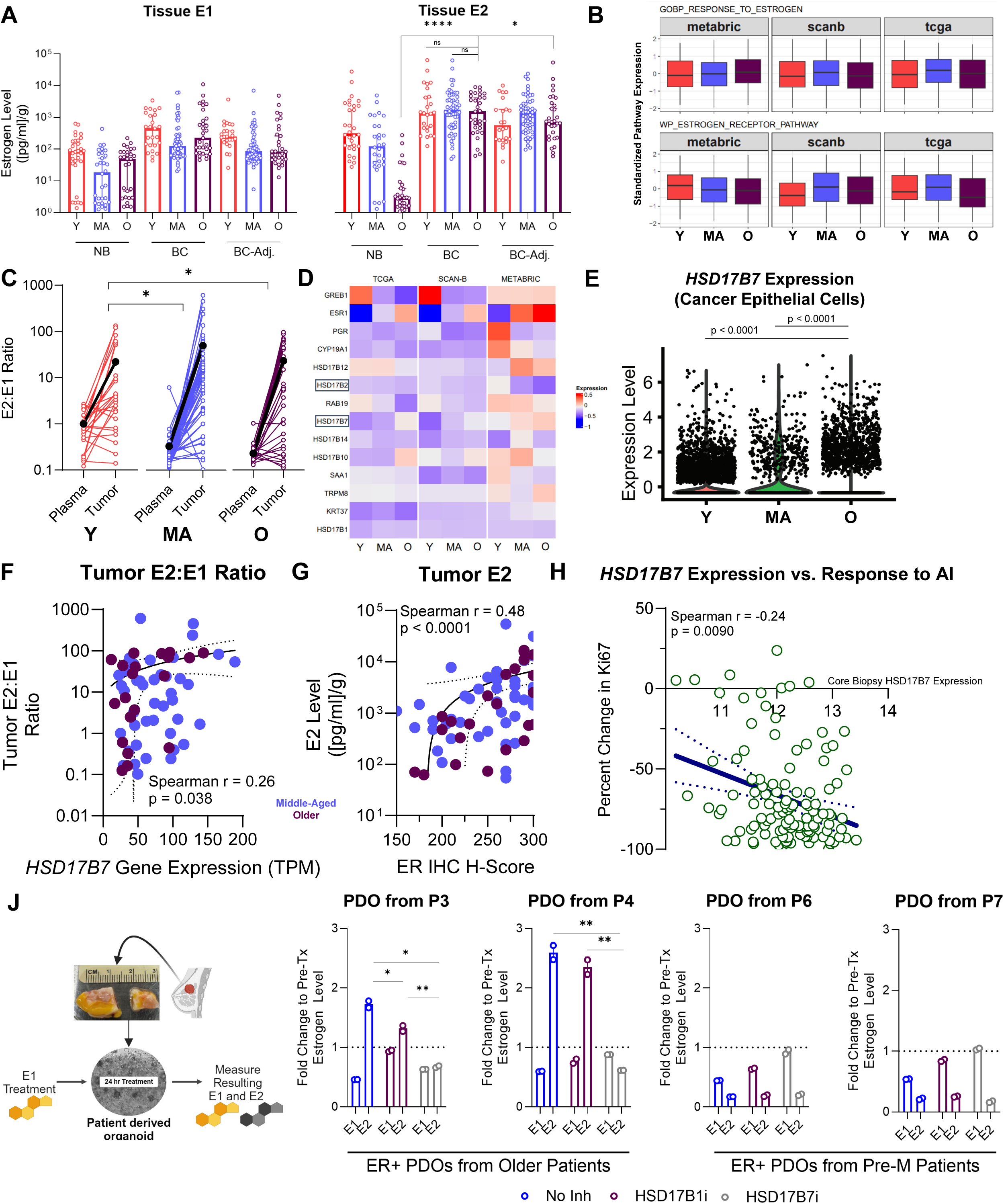
While E1 predominates in the circulating environment, age-related increases to *HSD17B7* expression fuel E1-to-E2 conversion in the breast, creating a local environment high in E2 and ER. (A) Tissue E1 and E2 measured across ages and across normal breast tissue, breast cancer tissue, and breast cancer-adjacent tissue. P-values can be found in data file S2. (B) With local E2 levels equivalent across age groups, downstream estrogen pathway activity show no trends by age when analyzed using the mutual information concordance analysis (MICA) pipeline. (C) E2:E1 ratio comparisons from the plasma to tumor across all ages in the ER+ breast cancer tissue. MA and O patients had significantly higher fold changes in plasma to tumor E2:E1 ratios compared to Y patients. * indicates p-value less than 0.05. Data corrected for multiple comparisons. (D) Profiling of a number of genes using the MICA pipeline to examine expression patterns across age groups using TCGA, METABRIC, and SCAN-B bulk RNA-seq datasets. Data show that only *HSD17B7* expression (increasing expression with increasing age) and HSD17B2 (decreasing expression with increasing age) were found to have consistent trends according to MICA. (E) *HSD17B7* levels were highest in older patients as corroborated by scRNA-seq data from ref. 68. (F) Scatter plot comparing *HSD17B7* expression levels (as measured using bulk RNA-seq) and measured tumor E2:E1 ratios, showing a positive correlation (Spearman r = 0.24, p = 0.03). Dot color indicates age group. (G) Scatter plot comparing ER IHC H-score and tumor levels of E2 ([pg/ml]/gram of tissue), showing a positive correlation (Spearman r = 0.45, p < 0.0001). (H) Using data from the POETIC trial (ref. 32), scatter plot compares baseline level of *HSD17B7* expression in the core biopsy against the percent change in Ki-67 after neoadjuvant endocrine therapy (Spearman r = -0.24, p = 0.009). (J) We treated patient-derived organoids (PDOs) with E1 and evaluated the degree to which that E1 was converted to E2 under the conditions of no inhibitor present, in the presence of an HSD17B1 inhibitor, or an HSD17B7 inhibitor. PDOs from Patients 3 and 4 (P3 and P4) were derived from older patients with ER+ breast cancer, whereas PDOs from Patients 6 and 7 (P6 and P7) were derived from younger, pre-menopausal patients with ER+ breast cancer (see Table S3 for full clinical characteristics of the PDOs used in this study).

Given the high E2 in the breast cancer TME, we hypothesized that ER activity should remain high in this setting. To test this, we examined a panel of estrogen-related pathways in early-stage ER+ breast cancer in the TCGA, METABRIC, and SCAN-B datasets. We used a novel statistical approach, mutual information concordance analysis (MICA), which evaluates consistent trends in pathway activity (or gene expression) within the three age groups across the three RNA-seq datasets (*31*). Concordant with the local E2 levels, downstream activity in estrogen-related pathways was not different across ages, indicating again that tumors in middle-aged and older patients have just as much estrogen signaling activity compared to younger patients (**Figs. 4B, S6A, and S6B**). When we compared plasma E2:E1 ratio against that found in the tumor tissue, we found that middle-aged and older patients had a significantly higher fold change increase compared to that found in younger patients (**Fig. 4C**), indicating that the tumors arising in the aged breast either synthesize local E2 de novo or have a mechanism to convert E1 to E2.

To decipher how E2 is synthesized in the breast, we examined a panel of enzymes involved in estrogen synthesis and conversion, again using the MICA method. Among all of the hydroxysteroid enzymes involved in E1 to E2 and E2 to E1 interconversion (schematic in **Fig. S7A**), *HSD17B7*, which is involved in E1 to E2 conversion, was found to be consistently increased and *HSD17B2*, involved in E2 to E1 conversion, consistently decreased with age across all three datasets, with no significant trends for other hydroxysteroid enzymes (**Fig. 4D**). This finding was restricted to tumor tissue, where *HSD17B7* expression was significantly higher when compared with normal-adjacent tissues (**Fig. S7B**). Aromatase (*CYP19A1*) was not significantly different across age groups. We found similar age-related increases in *HSD17B7* expression using ER+ breast cancer scRNA-seq data (*32*), specifically in the cancer epithelial cells (**Fig. 4E**), nominating HSD17B7 as a contributory enzyme to establishing an E2-rich TME in older patients.

In the same tissues in which estrogen disposition was measured, we performed bulk RNA-seq, which showed that *HSD17B7* expression was positively correlated with a number of estrogen-related pathways (**Fig. S8A**). Importantly, *HSD17B2* expression was negatively correlated with estrogen pathway activity (**Fig. S8A**). A derived signature of estrone and inflammatory (Estrone+TNF) activity (*30*) showed consistent decreases with age across TCGA (p = 0.049), METABRIC (p = 0.12), and SCAN-B (p = 0.0012) (**Fig. S8B**). However, analysis between *HSD17B7* expression and post-menopausal tumor E2:E1 ratio showed that as enzyme gene expression increased, so too did the balance of E2 to E1 in the TME (Spearman r = 0.26; p = 0.038) (**Fig. 4F**).

Consistent with prior reports, we find that *ESR1* gene expression increases precipitously with age (*8, 33*), a finding compounded with increasing hypomethylation and concomitant increases in ER protein (**Figs. 4D, S9A, and S9B**). Intriguingly, these changes are also seen with *ESR1* expression in normal-adjacent tissues from TCGA (**Fig. S9C**), suggesting that *ESR1* is likely sensitized to low plasma and breast E2 levels in post-menopausal women such that even the normal, non-cancerous breast harbors higher relative levels of ER than pre-menopausal counterparts. Uniquely within the aged TME, however, we observe that not only are there locally high E2 levels, but these tumors also retain high levels of ER positivity: tumors with the highest ER H-score also had high levels of tumor E2 (Spearman r = 0.48, p < 0.0001; **Fig. 4G**) whereas there was no association between ER H-score and measured E1 levels (**Fig. S9D**). The estrogen disposition of these tumors also enhances sensitivity to aromatase inhibition, where high levels of *HSD17B7* expression in the tumors of patients in the POETIC trial testing neoadjuvant aromatase inhibition (*34*) were associated with the largest percent change in Ki-67 (Spearman r = -0.24, p = 0.0090) (**Fig. 4H**).

Lastly, we utilized a panel of breast cancer cell lines and patient derived organoids (PDOs) to test their ability to convert E1 to E2. Treatment with physiologically-relevant levels of E1 (500 pg/ml) for 24 hours resulted in conversion of E1 to E2 in ER+ but not ER-human breast cancer cell lines, reproducing prior reports (*35*) (**Fig. S10A**). We then tested a series of PDOs from older patients with ER+ breast cancer as well as a PDO from a younger, pre-menopausal patient with ER+ breast cancer and an older patient with TNBC (full clinical characteristics of the PDOs used throughout this study can be found in **Table S3**). ER+ PDOs from older patients converted significantly higher levels of E1 to E2 than the PDO from a younger patient and the PDO from the older patient with TNBC (**Fig. S10B**). Given the age-related increase in HSD17B7 in ER+ breast cancer in older patients, we tested the role of HSD17B7 inhibition in preventing conversion of E1 to E2 in the PDOs using a specific small molecule inhibitor (Compound 10) (*36–39*). When we treated the PDOs with an HSD17B7 inhibitor (and used an inhibitor of HSD17B1, CC-156, as a control as this enzyme was not increased with age), the HSD17B7 inhibitor decreased E1 to E2 conversion in the ER+ PDOs from older patients with little effect on the other PDOs (**Fig. 4J**). The HSD17B1 inhibitor had little to no effect on E1 to E2 conversion. This result further solidifies the role of HSD17B7 in enhancing local E2 levels in the aged TME.

Overall, these results demonstrate that age-related breast tumors upregulate *HSD17B7* expression (and downregulate enzymes responsible for E2 inactivation) to create a unique local environment characterized by both high in E2 levels as well as high tumor ER levels.

### Aged breast TME exhibits features of chronic inflammation but is characterized by marked immune dysfunction

Given the chronic inflammatory phenotype seen in the plasma, we next investigated whether the breast microenvironment showed similar changes with age. While there was evidence of a modest chronic inflammatory response in normal breast tissue with age, older patients with ER+ breast cancer harbored high levels of a chemokine-enriched inflammatory tumor microenvironment compared to the tumor-adjacent tissue (**Figs. S11 and 5A**). This age-associated inflammatory environment was again most notably characterized by elevated levels of chemokines (though many factors, such as IL-6, IFN-y, IL-1β, and IL-8, did not change with age) indicating that the aged TME was chronically inflamed relative to the TME of both younger and middle-aged patients. We next examined the degree to which the inflamed TME shaped immune signaling in our patient cohort, where we measured local inflammatory factor levels and also performed bulk RNA-seq. We first segregated patients across age groups according to the levels of chemokine-defined inflammatory scores. Patients with the highest levels of chemokine inflammation in the TME harbored the lowest levels of immune pathway activation, indicating broad immune dysfunction in these patients, which was most prominent in older patients (**Fig. 5B**).

**Fig. 5:**
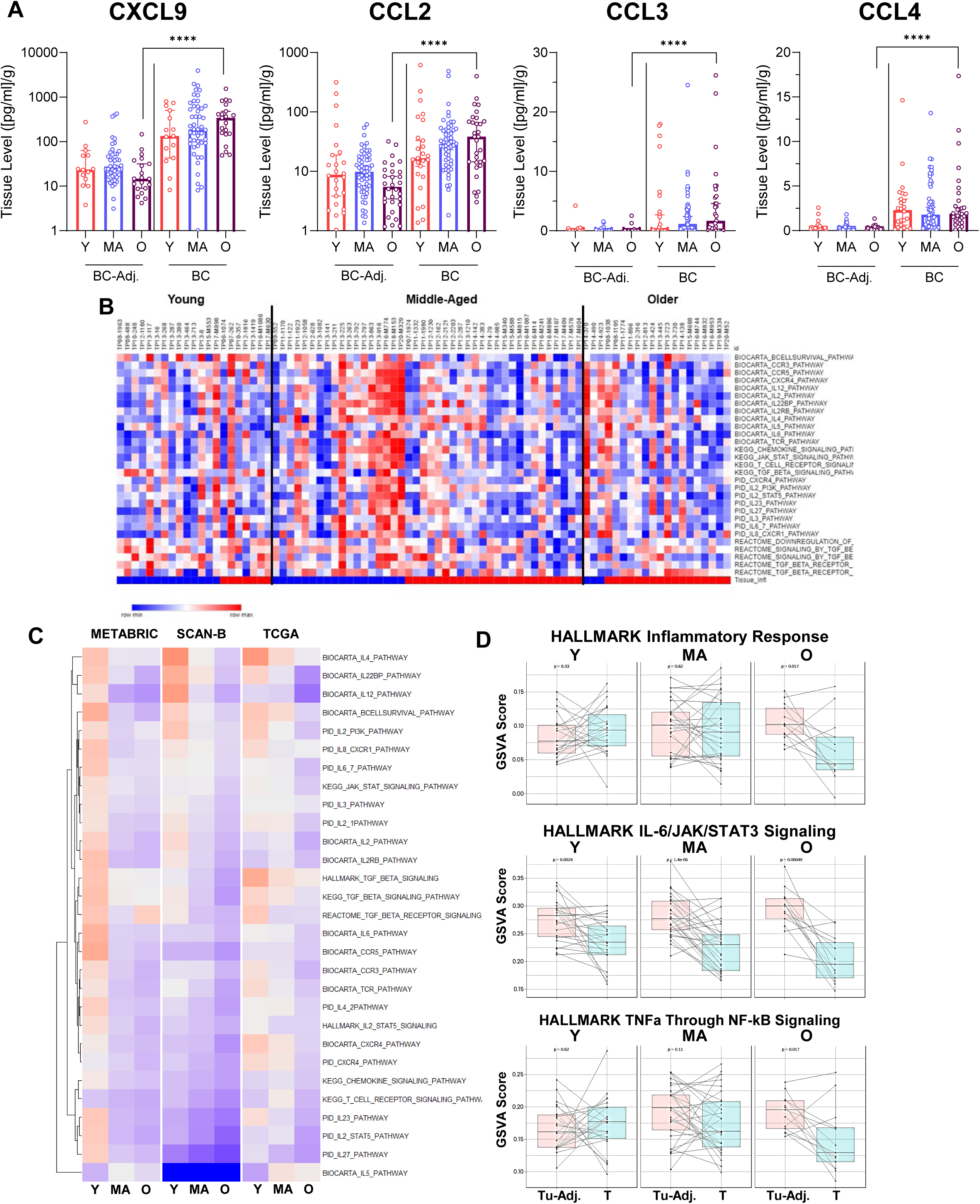
The aged breast TME recapitulates a chemokine-enriched chronic inflammation but is highly dysfunctional compared to younger and middle-aged patients. (A) Tumor lysates were analyzed using the same Luminex inflammatory panel. Similar to the circulating environment, the aged breast TME was characterized by high levels of chemokines, most notably CCL2. **** indicates p-value less than 0.0001. Data corrected for multiple comparisons. (B) Heatmap showing a series of inflammation-related pathways from MSigDB, indicating that across all three bulk RNA-seq datasets, older patients consistently had the lowest levels of pathway enrichment. (C) Shows comparisons between normal (tumor adjacent) and tumor tissue from TCGA according to enrichment of several HALLMARK pathways. Tumor tissue, but not normal tumor adjacent tissue, from older patients showed the lowest level of pathway enrichment. P-values indicated in figure. (D) Heatmap showing that for tumors that had the highest levels of chemokine inflammation scores (indicated by the “Tissue_Infl” row; calculated in red if a given patient had a chemokine score higher than the median across all patients) also had the lowest levels of inflammatory pathway enrichment, or the highest level of immune dysfunctionality.

Despite the inflamed nature of the TME, when we employed the MICA pipeline to profile all inflammation-related pathways across MSigDB, we observed broad age-related decreases in key inflammatory and immune activating pathways, with the lowest levels of pathway enrichment in older patients (**Fig. 5C**). Among the pathways differentially enriched, those related to reactive oxygen species and oxidative phosphorylation were consistently enriched in older versus both middle-aged and younger patients (**Fig. S12A**). However, pathways related to inflammatory signaling, such as TNFα through NF-κB, Hallmark Inflammatory Response, and IL-6-JAK/STAT, were consistently enriched in younger and middle-aged patients compared to older patients (**Fig. S12B**). Remarkably, in TCGA with matched tumor and tumor-adjacent tissues, while these pathways increased with age in the tumor-adjacent tissues, the tumors from older patients again exhibited the lowest scores (**Fig. 5D**).

Overall, while we observe a prominent inflammatory environment within the TME of older patients with ER+ breast cancer, these data suggest that the chronic inflammation is associated with immune dysfunction and suppression, indicating many of these tumors do not activate an immune response in a similar manner to that seen in both younger and middle-aged patients.

### ER+ tumors from older patients are characterized by immunosuppressive macrophages

Given the chemokine-enriched nature of both the circulating and local inflammatory environments, we hypothesized that macrophages may be a prominent component of and may shape the aged TME. Indeed, the proportion of cells from the monocytic lineage increased with age when examining deconvoluted data from bulk RNA-seq datasets (p-value for trend derived from MICA = 0.0012; **Fig. 6A**). We then investigated publicly available single-cell RNA-seq data (*32*), only including those tumors that were early-stage, untreated, ER+/HER2-(**Fig. S13A**). Across a broad range of immune-related pathways, including those that are typically found primarily in lymphocytes and found in myeloid cells, tumor-associated macrophages (TAMs) in older patients were amongst the most distinct from those in their younger (**Fig. 6B**) and middle-aged (**Fig. S13B**) counterparts. Notably, most immune activating pathways were lower in TAMs from older patients; only pathways such as hedgehog signaling, a pathway associated immunosuppressive macrophages, were enriched in TAMs in older patients (**Figs. 6B and S13C**).

**Fig. 6:**
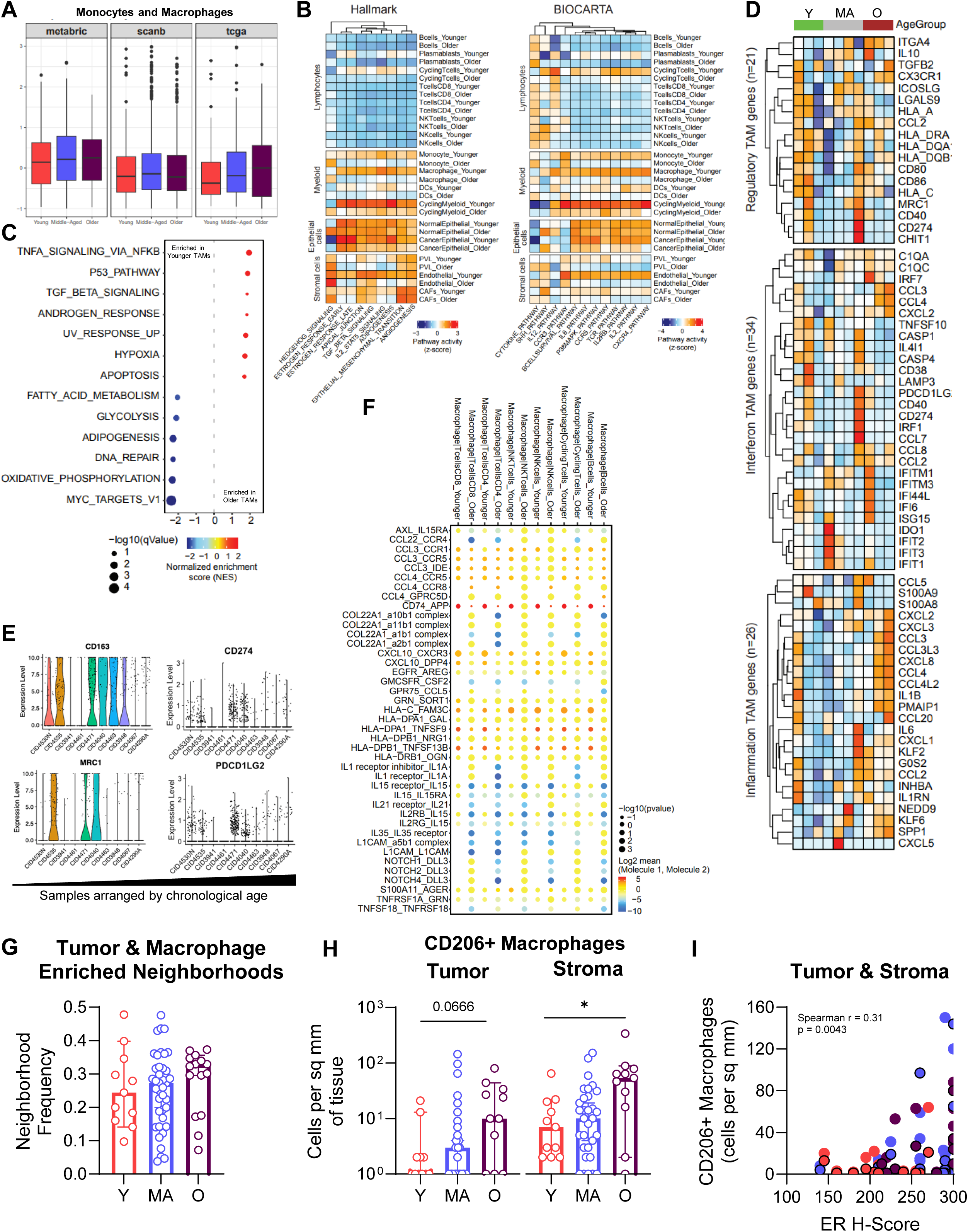
A chemokine-rich aged TME is associated with accumulation of immunosuppressive macrophages. (A) Using immune deconvoluted data from TCGA, METABRIC, and SCAN-B, the MICA pipeline identified that cells from the monocytic and macrophage lineage showed consistent increases against age. (B) Using publicly-available scRNA-seq data (ref. 68, data used in Figs. 6B through 6F; only used tumors that were treatment-naïve, ER+/HER2-), macrophages and monocytes were amongst the most transcriptomically different between age groups. (C) Differentially expressed gene (DEG) analysis with subsequent pathway analysis using the DEGs between macrophages from older patients and younger patients showed an enrichment of TNFa and TGFb signaling in older patients (indicated in red). (D) No consistent pattern was seen when subtyping the macrophages according the paradigm established in ref. 34, though we did observe that macrophages in older patients had consistently high expression of CCL2 and other chemokine-related genes. (E) Older patients had high levels of genes associated with M2-like macrophages, such as CD163 and MRC1 (CD206), as well as PD-L1 and PD-L2. (F) CellPhoneDB analysis shows that macrophages from older patients have less communication with adaptive immune cells compared to macrophages from younger patients. (G) Using previously-generated multispectral immunohistochemistry (mIHC) data from our lab (ref. 35), we reanalyzed the neighborhood analysis data from ER+/HER2-tumors according to age. There was a higher frequency of tumor and macrophage enriched neighborhoods in older patients. (H) Using the mIHC data from ref. 35, CD206+ macrophages were higher in both the tumor and stromal regions in older patients with ER+ breast cancer. * indicates p-value less than 0.05. (I) Using the mIHC data from ref. 35, we observed a positive association between tumor ER H-score and the amount of CD206+ macrophages present (Spearman r = 0.31, p = 0.0043). Dots colored according to age. Dots with black outline indicate cells from stromal regions while those without a border indicate cells from tumor regions.

We analyzed differentially expressed genes in TAMs from older patients compared to younger and middle-aged patients, which showed that while there was broad overlap in genes enriched in TAMs from younger and middle-aged patients, the TAMs from older patients seemed to harbor a unique set of genes (**Fig. S13D**).

Analysis on the DEGs indicated that pathways associated chronic inflammation, such as TNFα through NF-κB and TGFβ signaling, were enriched in TAMs from older patients compared to both younger and middle-aged patients (**Figs. 6C and S13E**). TNFα pathway activation was unexpected but likely related to DAMP activity and macrophage TLR signaling in the aged TME (**Fig. S13F**). Further subtyping of the TAMs according to the paradigm described by Ma et al (*40*) showed that while TAMs from older patients exhibited features of mapping to multiple TAM categories, these cells consistently had prominent markers associated establishment of an immunosuppressive TME, including chemokine signaling, *MRC1* (CD206) expression, and *CD274* (PD-L1) expression, which were collectively elevated in the middle-aged and older patients (**Figs. 6D and 6E**).

Hypothesizing that the immunosuppressive aged macrophages further shaped the local adaptive immune response, we looked at cell communication. TAMs from younger patients had higher degrees of immune activating communication to CD4 T cells, CD8 T cells, and NK cells, reflected in enhanced IL-2, IL-15, and IL-35 signaling, whereas the TAMs from older patients blunted the anti-tumor immune response (**Fig. 6F**).

To validate the scRNA-seq findings, we analyzed multispectral immunohistochemistry stained slides from a cohort of patients with ER+ breast cancer (*41*) with a focus on age-related changes. Compared to younger and middle-aged patients, while total macrophages did not change with age (in either the stroma or within the tumor; **Fig. S14**), the proportion of tumor and macrophage enriched neighborhoods increased in the TME in older patients (Fig. 6G), in part attributable to significant increases in tumor and stromal CD206+ TAMs with age (Fig. 6H) without any age-associated changes to the MHCII+ TAM infiltrate (**Fig. S14**). Importantly, the proportion of CD206+ TAMs was significantly associated with the degree of ER positivity in the tumor (Spearman r = 0.31; p = 0.0043). Collectively, these data suggest that CD206+ TAMs accumulate in older patients with ER+ breast cancer and drive an immunosuppressive program that promotes a blunted adaptive response.

### Aged TME promotes macrophage polarization to an immunosuppressive state

To mechanistically examine and validate the degree to which the aged TME, which is characterized not only by a chronically inflamed TME but also by high local E2 levels, enhances macrophage polarization, we used macrophages differentiated from monocytes (isolated from human donor PBMCs). To simulate the aged TME, we treated these macrophages with E2, CCL2, CXCL9, or the combination and quantified the staining intensity of the immunosuppressive markers CD206 and PD-L1. Treating with E2 alone significantly increased CD206 staining; the combination of all three agents promoted macrophage transition towards the CD206+/PD-L1+ state (**Fig. 7A**). Next, we prospectively collected a panel of ER+ PDOs from older patients with varying degrees of ER positivity and Ki-67 indices (**Table S3**). To capture the secreted contents of all cells in the aged TME, we generated conditioned media from zero passage, just-in-culture PDOs, and then incubated with monocyte-derived macrophages (n = 5) (**Figs. 7B and S13**). We measured baseline levels of inflammatory cytokines and chemokines as well as estrogens present in the PDO conditioned media. CCL2 and IL-6 were the most abundant inflammatory cytokines, while E2 was nearly 10-20 fold higher than E1. The conditioned media significantly increased both CD206 and PD-L1 positivity in macrophages, further confirming that the aged TME promotes the accumulation of immunosuppressive macrophages (**Fig. 7C**). Interestingly, when we again ran the same inflammatory panel on the media after the macrophages were polarized (containing secreted factors from the polarized macrophages), we observed a prominent secretory phenotype of IL-4, IL-10, IL-13, and TGFβ, functionally supporting the immunosuppressive nature of the macrophages (**Figs. 7D**). In particular, TGFβ significantly increased across the measured conditions (**Fig. 7D**).

**Fig. 7:**
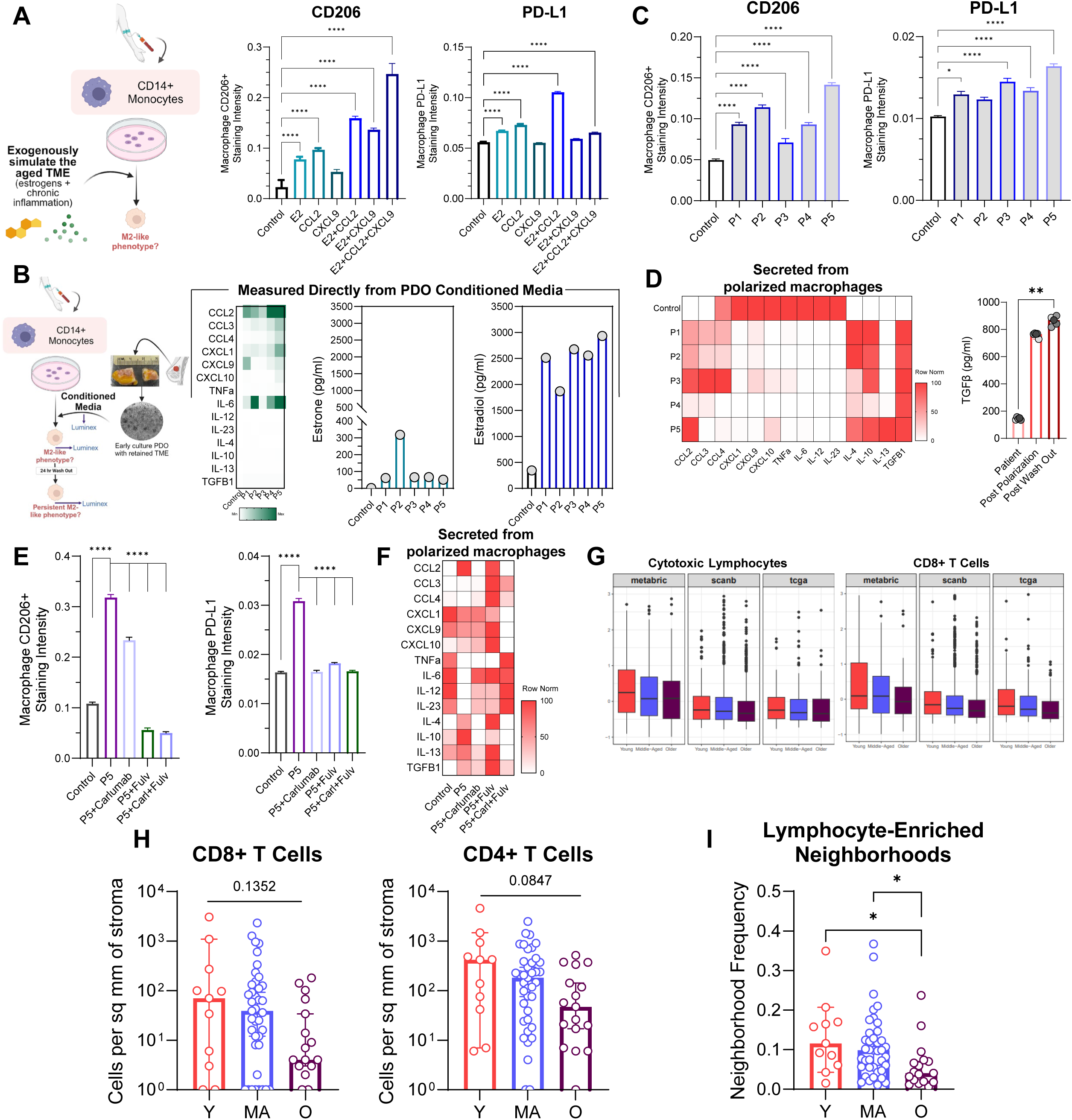
Aged TME polarizes macrophages to an immunosuppressive state that is targetable through the CCL2 axis or through endocrine therapies. (A) We first exogenously treated macrophages with components of the aged TME (including E2, CCL2, CXCL9 or the combination) and measured CD206 or PD-L1 staining intensity as assessed by immunofluorescence. Data show median with 95% CI. **** indicates p-value less than 0.0001. Data corrected for multiple comparisons. (B) We then extracted conditioned media from just-in-culture patient derived organoids (PDOs) from Patients 1-5 (P1-P5, full clinical characteristics of the collected PDOs can be found in Table S3). Heatmap shows non-normalized values of inflammatory markers present in the conditioned media or the control media (basal breast organoid media). We also measured E1 and E2 in the conditioned media, showing high levels of E2. (C) Adding the conditioned media from P1 to P5 significantly increased CD206 and PD-L1 staining intensity on macrophages. **** indicates p-value less than 0.0001. Data corrected for multiple comparisons. (D) After macrophage polarization, we again measured the media in an effort to measure what the macrophages were secreting. Heatmap shows row normalized values. TGFb was significantly increased after macrophage polarization and persisted after a 24 hour wash out period. ** indicates p-value less than 0.01. (E) Using the same experimental setup, we then treated macrophages with P5 conditioned media alone or in combination with carlumab (monoclonal antibody targeting CCL2) and/or fulvestrant (selective estrogen receptor degrader). **** indicates p-value less than 0.0001. Data corrected for multiple comparisons. (F) Heatmap showing inflammatory factors in the media after macrophage polarization or lack thereof in each of the conditions. (G) Using immune deconvoluted data from TCGA, METABRIC, and SCAN-B, the MICA pipeline identified that cytotoxic lymphocytes and CD8+ T cells showed consistent decreases with age. (H and I) Using the mIHC data from ref. 35, CD8+ and CD4+ T cells were lowest in the stromal regions from older patients with ER+ breast cancer; these tumors had significantly lower neighborhood frequencies characterized lymphocyte enrichment. * indicates p-value less than 0.05.

Given the degree of secretory immunosuppression from the PDO conditioned media, which contained both high levels of inflammatory factors and high levels of E2, we tested a number of inhibitors in an attempt to reduce the degree of macrophage polarization. When we again treated the macrophages with conditioned media from P5, macrophages again showed consistent polarization to a CD206+ state with increased secretion of CCL2, IL-4, IL-10, IL-13, and TGFβ (**Figs. 7E and 7F**). When we added either carlumab (anti-CCL2 monoclonal antibody) or fulvestrant (selective estrogen receptor degrader), macrophage polarization was significantly decreased, though to a far greater degree with fulvestrant (**Fig. 7E**). Interestingly, however, it was the combination treatment that both reduced IL-4, IL-10, IL-13, and TGFβ secretion and restored anti-tumor inflammatory factor secretion.

Lastly, with the immunosuppressive nature of the macrophages, we examined levels of lymphocyte infiltration in the tumors from older patients. Deconvolution of bulk RNA-seq data with analysis using the MICA pipeline showed consistent and significant age-related decreases to the CD8 T cell infiltrate in tumors from older patients along with a decrease in overall cytotoxic lymphocyte proportions (**Fig. 7G**). Similarly, pathway enrichment analysis corroboratively showed significant decrements in TCR signaling with age (**Fig. S14**). CD8 and CD4 T cell numbers were similarly decreased in our cohort of ER+ patients (analyzed using multiplexed IHC, **Fig. 7H**). Compared to younger and middle-aged patients, older patients with ER+ tumors exhibited significantly lower levels of lymphocyte-enriched neighborhoods (**Fig. 7I**), collectively suggesting an impaired downstream adaptive immune response associated with immunosuppressive macrophages.

## DISCUSSION

It is becoming increasingly recognized that ER+ breast cancer is an age-related disease, and due to the substantial prevalence of this patient population in the clinic, increased attention to both differences in clinical management and tumor etiology and biology is required (*7, 22, 42*). More so than younger patient counterparts, clinical management of older patients, typically defined as those over 70 years old, can be challenging in that tumor biology, life expectancy, competing risks of mortality, and patient preferences may all play a role in determining optimal therapies (*22, 42*). ER+ tumors in older patients tend to exhibit different clinical behavior, leading to recent changes in treatment paradigms in both the surgical setting (*43, 44*) as well as in the adjuvant setting (*45, 46*). Despite this, our understanding of the underlying tumor biology, immune biology, and role of the host environment and their contributions to tumor development, growth, and ultimately behavior is largely understudied. In this report, we use a novel aged rat model and a unique cohort of matched blood and breast tissue from both healthy donors without breast pathology and patients with ER+ breast cancer for a comprehensive systems biology analysis of the ecological makeup of aged breast cancers. We investigate the host (systemic) environment, how it shapes the aged breast tumor microenvironment, and how this hormonal and immune contexture fosters immune escape and tumor persistence. On the hormonal side, while paradoxical that so many ER+ breast cancers form later in life, when a woman is post-menopausal, the source of local estrogens in the post-menopausal breast has been an interest of the field for several decades

(*47*). However, because of the efficacy of aromatase inhibitors, interest into hormonal disposition in the breast largely waned. Despite this, identification and characterization of the role of local hydroxysteroid dehydrogenase activity, specifically in the aged breast, along with a potential strategy for inhibition, may rekindle interest in an emerging therapeutic avenue. On the inflammation side, there has been growing interest in better understanding how the host environment impacts the balance of pro- and anti-tumor immunity (*16, 18, 48*); however this necessitates a systems biology approach and thus few studies have comprehensively studied matched blood analytes, tumor microenvironment analytes, and tumor tissue sequencing.

We demonstrate that the aged environment in a rat model of ER+ breast cancer promoted accelerated yet indolent tumorigenesis. Our sequencing studies suggest that this effect was unlikely related to the accumulation of mutations from the carcinogen or from age alone and suggest a strong inflammatory component given the enrichment of chemokine-defined systemic inflammation and tumor-associated macrophage proportional increases in the aged rats. A finding that warrants future consideration is the perhaps paradoxical data that show despite cumulative immunosuppressive signals, tumors in older rats seem to grow slower, a phenotype also observed in aged breast cancers in humans. It is possible that the suppressed immune environment plays a more prominent role in initial immune escape and less so with regards to sustained tumor progression (*49, 50*). While this work implicates chronic inflammatory changes in the circulating environment and local immune contexture, tumor cell age-related changes certainly play a role in age-related tumorigenesis. Recent rodent work has elucidated potential tumor cell-driven changes, such as secretion of midkine, that reshape other mammary gland epithelial cells in the TME (*51, 52*). In addition to secretion of midkine, tumor cell expression of HSD17B7 may also create autocrine and paracrine signaling loops to increase local E2 levels as a fuel for growth. This dynamic nature of the age-related tumorigenesis certainly necessitates a holistic understanding of the interplay between systemic (host) factors and how these shape the local response to a growing tumor.

Importantly, our findings observed in the aged rat model were corroborated in the systemic and local environment in humans: while older patients harbored a chronic inflammatory circulating environment with an E1-predominant estrogen disposition, the aged local TME was uniquely defined by high tumor ER levels *and* high levels of local E2. Thus, the finding of HSD17B7-driven E2 accumulation in the aged TME is particularly engaging. Seminal studies several years ago suggested that local synthesis of E2 in the breast through aromatase activity was not the predominant modality driving intratumoral E2 levels (*53–55*). It was shown instead that hydroxysteroid dehydrogenase activity was the single most important determinant of in-breast E2 levels (*53*). Our work here suggests that for older patients, circulating E1 serves as an “estrogen pool” which is readily converted by the HSD17B7 enzyme, which was the only dehydrogenase to show age-related expression increases specifically in tumor tissue. The result is such that post-menopausal breast cancers have E2 levels similar to pre-menopausal breast cancers. Our findings here also minimize the potential role of local E1 activity in the setting of age-related breast cancer given the decrements in *HSD17B2* expression and far higher E2:E1 ratios in the breast tissue, even in older patients (*30, 56*). Interestingly, preceding the increase in local E2 is the fact that the normal breast likely increases *ESR1* expression (and thereby increasing ER expression) due to the low circulating and local levels of E2 (*57*). As the breast tumor forms, increases in HSD17B7 expression create a local environment that not only already has high levels of ER, but also then has high levels of local E2 to serve as a proliferation fuel, which may explain why these tumors are typically exquisitely sensitive to endocrine therapy in early stage disease settings. Even though aromatase inhibitor therapies provide sufficient coverage for preventing peripheral formation of both E1 and E2 from androgens, our study raises the possibility that local targeting of HSD17B7, a concept first proposed in 2011 (*47*), could potentially have a role in breast cancer therapy given the advent of newly-synthesized inhibitors specific to the estrogenic role of these enzymes (*38, 39*). Our data in an array of PDOs generated directly from patients with ER+ breast cancer show that this mechanism may be specific for older patients, since HSD17B7 inhibition had little activity in PDOs from younger patients. Given the systemic nature of aromatase inhibitors, side effects are numerous and often preclude adherence for the recommended five years. Local inhibition thus becomes attractive and may be achieved in a number of ways.

An equally important finding of this work is the chemokine-enriched chronic inflammation that defined both the systemic and local environments. It is increasingly well-established that older patients with breast cancer exhibit altered inflammatory profiles and immune changes with age (*58–60*). However, understanding how systemic inflammatory changes drive local changes has been challenging. In this study, chemokines, or chemotactic molecules, especially CCL2, seem to drive immunosuppressive macrophage accumulation in the aged breast TME, a finding corroborated by prior work in the setting of ER+ breast cancer (*61*). Interestingly, while we found high levels of chemokines within the TME, indicating a chronic inflamed environment, we also observed broad immune dysfunction as well. This CCL2 axis may play an important role in establishing early immune escape for these early-stage ER+ tumors through orchestration of impaired macrophage-T cell communication (*62–64*). While total macrophage levels did not change across age, we show that CD206+ macrophages, known for their pro-tumor functions, are highest in the aged TME. Critically, these CD206+ macrophages not only accumulate in response to high levels of CCL2, but also seem to further enhance local CCL2 levels event after polarization. Critically, targeting CCL2 not only reduced macrophage polarization to a CD206+/PD-L1+ state, but also reduces effector pro-tumor, immunosuppressive factors such as TGFb. These findings raise the idea that targeting CCL2 may not only work to treat the host environment, thereby reducing the systemic chronic inflammation, but may also serve to re-establish an intact anti-tumor immune response locally. In the setting of ER+ breast cancer, potential immune induction approaches targeting CCL2 may be a worthwhile and feasible strategy, either alone or in combination with endocrine therapy; data from this study suggest at least some synergism when targeting both CCL2 and ER, which not only reduce macrophage polarization, but also restore the balance between anti- and pro-tumor cytokines that communicate with adaptive immune cells.

An important implication of the work described here is the increasingly-recognized idea that macrophages (and potentially other immune cells) are sensitive to estrogens and endocrine therapies (*65, 66*). Here, we show that macrophages may serve as “hubs” of integration, integrating the unique signals of the TME such as the chronic inflammation (especially the chemokines) as well as the high local E2 levels. Indeed, macrophages seem to not only be sensitive to E2 but also polarized toward a CD206+ state when treated with E2 alone. Further, when we treated macrophages with conditioned media from just-in-culture PDOs from older patients with ER+ breast cancer, which were rich in CCL2 and E2, CD206 polarization was significantly higher and included potent downstream secretion of TGFβ. In this same system, macrophages were particularly responsive to treatment with fulvestrant, which has been shown in other settings, such as melanoma, to decrease immunosuppressive properties of macrophages, thereby restoring anti-tumor immunity (*67*). Ongoing observational trials are now seeking to better understand the role endocrine therapies play in modulating systemic and local immunity (*68*) and will be important for better understanding how macrophages may aid anti-tumor responses after endocrine therapy.

In summary, the work presented here provides new insights into the unique systemic and local hormonal and inflammatory milieu that creates a permissive environment for ER+ breast cancer development and growth. This study uncovers a number of new therapeutic avenues, such as targeting local HSD17B7 enzymes or CCL2 to dampen the immunosuppressive features of the aged environment, that will require in vivo and early phase clinical trials for evaluation. These data provide strong rationale for age-specific therapeutic modalities for future in vivo studies and human trials.

## MATERIALS AND METHODS

### Study design

The objective of this study was to gain insight on why ER+ breast cancer is an age-related disease. While prior work has focused on age-related cancer cell-intrinsic changes, we hypothesized that the environment, collectively encompassing both the systemic and local environments, contributes to breast tumor development and growth in older patients. To examine this, we used an immunocompetent rat model of ER+ breast cancer, a cohort of clinical specimens from patients with ER+ breast cancer and from donors in the Komen Tissue Bank without known breast disease, publicly available scRNA-seq and bulk RNA-seq datasets, and a prospectively collected panel of tumor organoids from older patients with ER+ breast cancer.

All studies with human tissue were conducted in accordance with protocols approved by the Institutional Review Boards (IRB) of the University of Pittsburgh (STUDY20040297), the University of Pittsburgh Medical Center (IRB0502025 / HCC protocol 04-162), and Indiana University. Informed consent from each patient or donor was obtained before tissue collection and banking.

Studies using archived frozen and FFPE human specimens were coded and processed and analyzed in a blinded fashion. Studies using prospectively collected patient derived organoids were not blinded; however, data was analyzed using imaging methodologies that limited bias (as described in the sections below). Sample size calculations were not performed for human breast tissues studies; the sample size was determined by the number of available specimens with matched blood, tumor tissue, and tumor-adjacent tissue. In the rat studies, power calculations to determine sample size were calculated based on differences in expected tumor incidence rates in the young and old rat groups. Blind independent review of tumor histology, ER staining, and Ki-67 was performed by a trained animal pathologist; a minimum of four tumor areas were assessed.

Any sample exclusion is as described below or in the figure legend. All in vitro studies were replicated a minimum of two times as described in the figure legends.

### Rat experiments

All animals were housed in The Ford / Assembly Research Building at the University of Pittsburgh in an approved animal facility. All animal experiments were performed following protocols 20108061 and 23103979, which were both approved by the University of Pittsburgh Institutional Animal Care and Use Committee. Virgin Fischer 344 (F344) rats aged 6-8 weeks old were purchased from Envigo and served as the “young” rats in the study. Aged F344 rats were obtained from the National Institutes of Aging (NIA) Rodent Colonies at 19-21 months old and served as the “older” rats in the study. All rats were housed in, ventilated autoclaved caging systems (Allentown Caging) on hard chip bedding, had 12hr/12hr light-dark cycles, were provided with hyperchlorinated RO water, and were fed ad libitum rodent chow (Purina). Rats lived in the facility for at least two months prior to any experimental treatment to allow for equilibration.

In order to better understand the ovulatory status of the rats in each age group, vaginal cytology was performed on a cohort of rats not treated with DMBA or MPA and was performed for 14 consecutive days for 6 young and 6 older rats using a previously established protocol (*69*). Cytology specimens were dried overnight and stained using Diff-Quik staining protocol. Final cytology reads of the estrous cycle were done so in a blinded fashion with respect to the animal age.

### DMBA/MPA model

To investigate the effect of age on tumor incidence and growth, we used a chemical carcinogen model. All rats were surgically implanted with a 50mg MPA pellet in a subcutaneous fashion (90 day continuous release; Innovative Research of America, #NP-161). One week following pellet implant, rats were dosed with a single oral gavage of DMBA (65mg/kg; Thermo Scientific Chemicals, #408181000) dissolved in sesame oil (Thermo Scientific Chemicals, #241002500). All rats recovered fully from pellet implant surgery and tolerated the DMBA dose.

### Tumor growth analysis and tissue harvesting

Mammary tumor development and growth were monitored biweekly by palpating all six mammary gland pairs. Once a tumor formed, we assessed growth dynamics using a volumetric calculation of tumor growth ([L*W*W]/2). The experimental endpoint was reached once the tumor reached 3000mm^3^ or when the rat showed signs of clinical decline. Tumors were harvested and stored as cryopreserved specimens, were flash frozen, or further processed for paraffin embedding (10% formalin overnight, stored thereafter in 70% ethanol). All tumors were stained for ER (abcam, #ab32063, 1:10,000 dilution) and Ki-67 (Novus Biologicals, NB600-1252, 1:100 dilution).

All tumors were histologically confirmed by an experienced animal pathologist. All subsequent grading (mitoses per high power field, percent of tumor cells positive for ER, percent of tumor cells positive for Ki-67, and stromal TILs) was done so in a blinded fashion to the age of the animal and was a calculated average of four different areas of each tumor specimen.

### Rat tumor sequencing and computational analysis

A group of six flash-frozen tumors (all three tumors from younger rats and three tumors with adequate tissue from older rats) were processed and underwent single-nuclei RNA-sequencing, low-pass whole genome sequencing, whole exome sequencing, and bulk RNA-sequencing.

#### Single-nuclei RNA-sequencing

We performed 3’v3.1 Nuclei Isolation and GEX library preparation with the 10X Genomics Platform. Snap frozen tissues are dissociated with the 10X Genomics Nuclei Isolation Kit. After dissociation, nuclei suspension concentration and nuclei integrity are assessed using AOPI on the Nexcelom Cellometer Auto 2000. Libraries are then prepared according to the Chromium Next GEM Single Cell 3ʹ Reagent Kits v3.1 (Dual Index) protocol using the 10X Genomics Chromium iX Instrument. The resulting full-length cDNA is checked for size and concentration on the Agilent Tapestation 4150 before proceeding with gene expression library preparation and sample indexing. Final libraries are checked for size on the Agilent Tapestation and concentration using the Invitrogen Qubit 2.0. Libraries are pooled and sequenced on the Illumina NovaSeq 6000 Instrument.

#### Whole exome sequencing

Genomic DNA samples were quantified using Qubit 2.0 Fluorometer (ThermoFisher Scientific, Waltham, MA, USA). Enrichment probes were designed against the region of interest and synthesized through Agilent Technologies–SureSelectXT CD Rat All Exon. Library preparation was performed according to the manufacturer’s guidelines. Briefly, the genomic DNA was fragmented by acoustic shearing with a Covaris S220 instrument. Fragmented DNAs were cleaned up and end repaired, as well as adenylated at the 3’ends. Adapters were ligated to the DNA fragments, and adapter-ligated DNA fragments were enriched with limited cycle PCR. Adapter-ligated DNA fragments were validated using Agilent TapeStation (Agilent Technologies, Palo Alto, CA, USA), and quantified using Qubit 2.0 Fluorometer. Adapter-ligated DNA fragments were hybridized with biotinylated baits. The hybrid DNAs were captured by streptavidin-coated binding beads. After extensive wash, the captured DNAs were amplified and indexed with Illumina indexing primers. Post-captured DNA libraries were validated using Agilent TapeStation (Agilent, Santa Clara, CA, USA) and quantified using Qubit 2.0 Fluorometer and Real-Time PCR (KAPA Biosystems, Wilmington, MA, USA). The sequencing libraries were multiplexed and clustered onto a flowcell on the Illumina NovaSeq instrument according to manufacturer’s instructions. The samples were sequenced using a 2×150bp Paired End (PE) configuration. Image analysis and base calling were conducted by the NovaSeq Control Software (NCS). Raw sequence data (.bcl files) generated from Illumina NovaSeq was converted into fastq files and demultiplexed using Illumina bcl2fastq 2.20 software. One mis-match was allowed for index sequence identification.

Sequencing results were aligned and analyzed using version 3.4.0 of the NF-Core Sarek Pipeline, using Nextflow version 23.10.1, Singularity 3.9.6. Pipeline settings used the Singularity profile, Rnor_6.0 Rat genome, CNVKit and Mutect2 tools, skipped base_recalibrator, WES-only mode with a minimal SLURM config file for the University of Pittsburgh Center for Research Computing High Throughput Computing Cluster. Mutect2.vcf.gz outputs were then annotated using VEP 95. High impact and Moderate Impact mutations as defined by Ensembl were then extracted for analysis. Rat genes were made comparable to human homologs using PyBioMart, and then filtered down to a list of pan-cancer and breast cancer genes (See supplementary information), in cases where multiple gene mutations were found in the same gene, ties were broken by prioritizing greatest impact followed by frequency. Results were then plotted in an Oncoplot. Tumor Mutational Burden was calculated using the whole exome length of the rat genome of 71Mb. K-means clustering was done by selecting all genes affected by moderate and high impact mutations and binarized into a corresponding matrix. Predicted labels were then evaluated to see if the two populations of tumor were distinctly separable based on mutational characteristics. Mutational Signature Analysis was done using SigProfilerAssignment, with context 96 and genome Rnor_6.0. CNVKit analysis was run in tumor-only mode, without normal control due to a sample quality issue that affected CNVKit output.

#### Bulk RNA-sequencing

Total RNA was extracted from fresh frozen tissue samples using Qiagen RNeasy Plus Universal mini kit following manufacturer’s instructions (Qiagen, Hilden, Germany). RNA samples were quantified using Qubit 2.0 Fluorometer (ThermoFisher Scientific, Waltham, MA, USA) and RNA integrity was checked with 4200 TapeStation (Agilent Technologies, Palo Alto, CA, USA). ERCC Ex-fold RNA reagent (Cat: 4456739) from ThermoFisher Scientific, was added to normalized total RNA prior to library preparation following manufacturer’s protocol. The next steps included performing rRNA depletion using QIAseq® FastSelect™−rRNA HMR kit (Qiagen, Germantown, MD, USA), which was conducted following the manufacturer’s protocol. RNA sequencing libraries were constructed with the NEBNext Ultra II RNA Library Preparation Kit for Illumina by following the manufacturer’s recommendations. Briefly, enriched RNAs are fragmented for 15 minutes at 94°C. First strand and second strand cDNA are subsequently synthesized. cDNA fragments are end repaired and adenylated at 3’ends, and universal adapters are ligated to cDNA fragments, followed by index addition and library enrichment with limited cycle PCR. Sequencing libraries were validated using the Agilent Tapestation 4200 (Agilent Technologies, Palo Alto, CA, USA), and quantified using Qubit 2.0 Fluorometer (ThermoFisher Scientific, Waltham, MA, USA) as well as by quantitative PCR (KAPA Biosystems, Wilmington, MA, USA). The sequencing libraries were multiplexed and clustered onto a flowcell on the Illumina NovaSeq instrument according to manufacturer’s instructions. The samples were sequenced using a 2×150bp Paired End (PE) configuration. Image analysis and base calling were conducted by the NovaSeq Control Software (NCS). Raw sequence data (.bcl files) generated from Illumina NovaSeq was converted into fastq files and de-multiplexed using Illumina bcl2fastq 2.20 software. One mis-match was allowed for index sequence identification. We used FastQC/0.11.7 to check the quality of reads followed by Trimmomatic/0.38 to remove the adapter contents. High quality reads(Q>30) were used for alignment using Star/2.7.9a followed by Picard/2.18.12 to mark duplicate and then run HTSeq/0.13.5 to count the reads and then DESeq2 to identify differentially expressed genes.

### Patient cohorts and clinical specimens

#### Patients with ER+/HER2-breast cancer from PBC

To assemble a cohort of patients with ER+/HER2-breast cancer across age groups, we obtained archived specimens from patients that had matched tumor tissue, tumor-adjacent tissue, and blood from the University of Pittsburgh’s Pitt Biospecimen Core (PBC). No patients included in this cohort received neoadjuvant systemic therapy and were considered treatment naïve at the time of tissue and blood collection. Tumor-adjacent tissue was taken approximately 3-4mm away from the tumor in lumpectomy cases and 5-6mm away from the tumor in mastectomy cases. All patients signed informed consent for their specimens to be used in research applications. We collected patient and tumor characteristics associated with the specimens. All patients were diagnosed with ER+/HER2-early-stage breast cancer. We obtained specimens across young (35-45yo), middle-aged (55-69yo), and older (≥ 70yo) age groups.

#### Specimens from donors participating in the Komen Tissue Bank

To assemble an age-matched control cohort of women without clinical breast disease, we obtained fresh frozen normal breast tissue cores and matched serum from the Susan G. Komen Tissue Bank at IU Comprehensive Cancer Center. Details of the KTB project (http://komentissuebank.iu.edu/) have been previously described (*70, 71*). At the time of donation, participants provided written informed consent and were enrolled under a protocol approved by the IRB at Indiana University. We again obtained specimens across young (35-45yo), middle-aged (55-69yo), and older (≥ 70yo) age groups. Relevant exposures, such as age at menarche and hormone replacement therapy use, are recorded in Supplementary Table S2.

### Quantification of inflammatory factors

To analyze the inflammatory burden in the human serum, tissue microenvironment, rat serum, and PDO conditioned media, we utilized the Luminex multiplex system. In brief, beads of defined spectral properties conjugated to analyte specific capture antibodies and samples (including standards of known analyte concentration, control specimens, and unknowns) were pipetted into the wells of a filter bottom microplate and incubated. During this first incubation, analytes bind to the capture antibodies on the beads. After washing the beads, analyte-specific biotinylated detector antibodies were added and incubated with the beads. During this second incubation, the analyte-specific biotinylated detector antibodies recognize their epitopes and bind to the appropriate immobilized analytes. After incubation period of biotinylated detector antibodies, streptavidin conjugated to the fluorescent protein, R-Phycoerythrin (Streptavidin-RPE), was added (without washing) and incubated. During this final incubation, the Streptavidin-RPE binds to the biotinylated detector antibodies associated with the immune complexes on the beads, forming a four-member solid phase sandwich. After washing to remove unbound Streptavidin-RPE and biotinylated detector antibodies, the beads were analyzed with the Bio-Plex 200™ instrument using Bio-Plex Manager 6.2 software.

For all human studies (including human serum and tissue lysates), we ran a 22-plex panel consisting of GM-CSF, CXCL1, IFNy, IL-1b, IL-6, IL-8, IL-10, IL-17A, CCL2, CCL3, CCL4, TNFa, G-CSF, M-CSF, IL-1a, IL-2, IL-5, IL-13, IL-15, IL-18, CXCL10, and CXCL9. For all rat serum studies, we ran an 18-plex panel consisting of IFNy, IL-1b, CXCL1, IL-6, IL-10, IL-17, G-CSF, IL-18, IL-5, CCL3, IL-1a, CCL2, IL-2, CXCL10, IL-13, GM-CSF, TNFa, and IL-15. Lastly, for the human PDO conditioned media analysis, we ran a 15-plex consisting of CCL2, CCL3, CCL4, CXCL1, CXCL9, CXCL10, TNFa, IL-6, IL-12, IL-23, IL-4, IL-10, IL-13, and TGFb.

### Mass spectrometry for estrogen (E1 and E2) measurements

A previously validated LC-MS/MS assay for the quantification of estradiol and estrone in human serum was adapted and used for this study (*72*). Briefly, a LC-MS/MS system consisting of a Thermo Scientific Vanquish UPLC and TSQ Altis that was equipped with a heated ESI (HESI) source was used while the SRM transitions used for quantitation were m/z 504.3 →171.1 for estrone and m/z 506.2 →171.1 for estradiol. 500µL of serum and 25µL of internal standard underwent liquid-liquid extraction with 3mL of N-butyl chloride. The sample was briefly vortexed and centrifuged at 3,000 x g for 5 minutes before the organic layer was transferred and evaporated under nitrogen. The samples were derivatized using 50µL of 50 mM sodium bicarbonate buffer in addition to 50µL of dansyl chloride (1 mg/mL) and was then being heated for 3 minutes at 60°C. The samples were transferred to LC vials and 7.5µL was injected into the LC-MS system. Chromatographic separation of the samples was accomplished with a Waters Acquity BEH C18 (2.1 x 150 mm, 1.7 µm) column with a gradient elution using 0.1% formic acid (A) and acetonitrile (B) at a flow rate of 0.3 mL/min. The total runtime was 6.5 minutes.

### Bulk RNA-seq of ER+ breast cancer and computational analysis

We used the standard procedure of RNeasy kit to extract total RNA from ER+ breast cancer samples of 83 tumor or 85 tumor-adjacent normal tissue obtained from Pitt Biospecimen Core (PBC) and grouped them by chronological age. The NovaSeq 6000 library for DNA sequencing was prepared using TruSeq Stranded mRNA Library Prep Kit (Illumina) following the protocol provided by the manufacturer. Library size and quality were checked using Fragment Analyzer (Agilent) and fluorescent quantification on the Infinite F Nano + (Tecan). The libraries were normalized and pooled, and then sequenced using the NovaSeq 6000 Platform (Illumina) to an average of 25M 101 PE reads. Sequencing data was given as raw data with a Phred Q30 score of 80 or better. For processing, we used Rsubread (Bioconductor release 2.16) (*73*) to align sequence reads to reference genome and used edgeR (*74*) and limma (*75*) R packages (Bioconductor release 4.0 and 3.58 respectively) to normalize gene expression level to log2 transcripts per million (TPM) (*76*). We aligned sequence reads to GRCh38 release 14 human genome reference sequence and mapped the aligned sequences to Entrez Genes. After normalization, we removed genes of which expression level is zero across all samples to get 39,404 genes for further pathway analysis.

Our investigation into pathway activities involved the utilization of two distinct R packages: PROGENy (version 1.22) (*77*) and GSVA (version 1.48.3) (*78*) with default parameters. PROGENy is a resource that leverages a large compilation of signaling perturbation experiments for a common core of pathway-responsive genes.

GSVA provides heightened sensitivity for detecting subtle changes in pathway activity across a sample population. In our GSVA analysis, we employed gene set collections associated with the estrogen pathway. These gene sets were sourced from hallmark, reactome, wikipathways, and gene ontology biological pathways. These two tools allowed us to infer the activities of biological pathways. We calculated Spearman correlation coefficients between the inferred pathway activities in single sample and expression of genes involved in estrogen conversion.

### Publicly available data sets used

#### scRNA-seq of treatment-naïve, ER+ breast cancer

We compiled single-cell RNA sequencing (scRNA-seq) datasets from ten ER+ breast tumor samples, focusing exclusively on cells from pre-treatment patients. The processed scRNA-seq data, originally published by Wu et al. (*79*) were obtained from the NCBI Gene Expression Omnibus (GEO) under accession number GSE176078. To analyze the data, we imported the matrix.mtx, features.tsv, and barcodes.tsv files were imported into a Seurat object using the Read10X function in the Seurat R package (version 5.0.3) (*80*). Our cell selection criteria included expression of fewer than 6,000 genes, with a minimum of 200 expressed genes and 400 UMI counts. Additionally, we filtered out cells with more than 15% of reads mapped to mitochondrial gene expression. To address technical variations and batch effects, we applied SCTransform to each Seurat object (*81*). SCTransform is a technique designed to mitigate technical variations and alleviate batch effects in scRNA-seq datasets. Principal component analysis (PCA) was performed on the filtered feature-by-barcode matrix, and Uniform Manifold Approximation and Projection (UMAP) embeddings (*82*) were based on the first 30 principal components. Overall, our processed dataset comprised 29,733 genes across 30,959 cells. For cell type annotation, we followed the major and subset cell types provided by Wu et al.

We employed CellPhoneDB (version 4) (*83*) to investigate potential ligand-receptor interactions between macrophages and other immune cells. Specifically, we analyzed raw count matrices separately for the younger and older groups. For each group, we calculated the mean average expression levels of interacting ligands in the sender population and interacting receptors in the receiver population. To assess the statistical significance of each interaction score, we performed a one-sided Wilcoxon signed-rank test.

#### Bulk RNA-seq data sets: TCGA, METABRIC, SCAN-B

We utilized the TCGA (*84*), METABRIC (*85*), and SCAN-B (*86, 87*) datasets for bulk RNA-seq analysis. Within each dataset, we only selected ER+/HER2-tumors within the designated age groups defined in this study. To identify biomarkers with consistent expression patterns across various age groups in TCGA, METABRIC, and SCAN-B datasets, we utilized the Mutual Information Concordance Analysis (MICA) framework (*31*). MICA, a novel in-house statistical method, employs mutual information to detect biomarkers with concordant multi-class (for example, by age group) expression patterns across multiple studies (for example, TCGA, METABRIC, and SCAN-B). Initially, it performs a global test to evaluate overall concordance, followed by a post hoc analysis to pinpoint specific studies contributing to this concordance. This approach ensures robust detection of biomarkers exhibiting consistent expression across multiple datasets.

#### Multispectral IHC cohort of ER+/HER2-IDC and ILC

To examine changes in immune cells and cellular neighborhoods across age groups, we utilized a previously developed cohort of patients with ER+/HER2-IDC and ILC (*41*). We re-analyzed the data according the age groups defined in this study; further details on antibodies and staining, image acquisition and cell quantification, and cellular neighborhood analysis can be accessed through the work by Onkar and colleagues (*41*).

### Patient-derived organoid generation, culture, and conditioned media

Patient-derived organoids were prospectively collected from consecutive patients over 70 years old with ER+/HER2-tumors who signed consent for specimens to be involved in research. Primary human breast cancer tissue was obtained by the Pitt Biospecimen Core from patients consented under the Breast Disease Research Repository (BDRR): Tissue and Bodily Fluid and Medical Information Acquisition Protocol (HCC 04-162). Patient-derived organoids were generated from this tissue in accordance with IRB STUDY22030183 by the Institute for Precision Medicine according to established protocol (*88*). Breast Organoid Medium (BOM) consisted of Advanced DMEM:F12 (+ 1X Glutamax, 10mM HEPES, 1X antimicrobial/antimycotic), 1nM β-estradiol, 500nM A83-01, 5ng/ml EGF, 250ng/ml R-Spondin, 100ng/ml Noggin, 5nM Heregulinβ-1, 5ng/ml FGF7, 20ng/ml FGF10, 500nM SB-202190, 5µM Y-27632, 1.25mM N-Acetyl-L-cysteine, 5mM Nicotinamide, 1X B27 Supplement, 50µg/ml Primocin. Briefly, tumors were digested with 2mg/ml collagenase (Sigma C9407) on a rotator, sheared, filtered at 100µm, and embedded in 40µl domes of Cultrex growth factor reduced BME type 2 (Fisher 35-330-1002) in 24-well non-treated plates (Fisher 12-566-82). At passage zero, as soon as the organoids are established in culture, 600µl BOM was added to each well and conditioned medium from organoids was collected after 2-3 days.

### Patient-derived organoid and cell line experiments testing for conversion from E1 to E2

PDOs from older patients with ER+ breast cancer, from a younger, pre-menopausal patient with ER+ breast, and from an older patient with TNBC were seeded approximately 10 days prior to experiment commencement. We note that all PDOs were collected from patients that did not receive any neoadjuvant systemic therapy. Twenty-fours hours prior to treatment with E1, a wash out of the PDOs was performed and breast organoid media without E2 was added to the culture. On the day of experiment start, PDOs were then treated in duplicate with 1.5nM E1 without E2 for 24 hours. After 24 hours, the media was collected and sent for mass spectrometry analysis as described above. We subsequently employed the same system to test two inhibitors to reduce E1 to E2 conversion, including Compound 10 (HSD17B7 inhibitor, treatment at 1µM for 24 hours) (*37*) and CC-156 (HSD17B1 inhibitor, treatment at 1µM for 24 hours) (*89*).

### Macrophage polarization experiments

PBMCs were isolated from healthy donors using Ficoll (Cytiva, #17-5446-52) and SepMate-50 tubes (STEMCELL, #85460) for density gradient centrifugation according to the manufacturer’s procedure. Monocytes were isolated from PBMCs (pan monocyte isolation kit Miltenyi, #130-096-537) prior seeding in a 96 well plate and differentiation to macrophages with 5ng/ml human M-CSF for two days. Patient-derived organoid conditioned medium (PDO CM) was collected and diluted at 1:1 ratio with fresh RPMI full medium. After washing out the M-CSF with PBS, diluted PDO CM was added to the monocyte-derived macrophages for polarization. After three days, samples are fixed with 4% PFA, permeabilized with 0.1% Triton X-100 and blocked for immunofluorescent (IF) staining. IF staining images were analyzed with CellProfiler (*90*) to determine the mean intensity of macrophage polarization markers of each cell. We quantified the expression levels of CD206 (Biolegend, #321102) and PD-L1 (Biolegend, #329701) in macrophages. To test if we could block macrophage polarization, we added carlumab (an anti-CCL2 antibody; MedChemExpress HY-P99188), fulvestrant (a selective estrogen receptor degrader; ICI 182780), and/or an IgG control antibody to the macrophage-PDO CM system.

### Statistical analysis

To assess differences in baseline characteristics in the human tissue cohorts, Fisher exact testing was performed for categorical variables and Mann Whitney testing was performed for continuous variables. Correlation analyses through the manuscript were assessed by nonparametric Spearman’s rank testing. A multivariable linear model was used to test the association between plasma and tissue inflammatory factors and chronological age. Two-tailed Mann Whitney testing was performed for all pairwise testing; ANOVA with a correction for multiple comparisons was performed for all other multi-group comparisons. Specific statistical tests for each analysis are stated in the figure legends.

## Acknowledgements

We thank all patients who consented for tissue and blood donation through Magee Women’s Hospital Women’s Cancer Research Center. Samples from the Susan G. Komen Tissue Bank at the IU Simon Cancer Center were also used in this study, and we thank contributors, including Indiana University who collected samples used in this study, as well as donors and their families, whose help and participation made this work possible. We thank Mitch Dowsett for reviewing early versions of the manuscript. We thank the Pitt Biospecimen Core for banking and distribution of de-identified biosamples and Sharon Winters and Vonda Mazzarella of the UPMC Hillman Cancer Registry for providing clinical data as honest brokers. Work performed in the UPMC Cancer Proteomics Facility (Luminex Core Laboratory) and services and instruments used in this project were graciously supported, in part, by the University of Pittsburgh. This project used the service of the University of Pittsburgh Small Molecule Biomarker Core facility, which was supported, in part, by the University of Pittsburgh Office of the Senior Vice Chancellor, Health Sciences, and the NIH S10RR023461 (to TDN) and S10OD028540 (to TDN). Work performed in the UPMC Hillman Cancer Center Tissue and Research Pathology/Pitt Biospecimen Core and services and instruments used in this project were supported, in part, by the University of Pittsburgh, the Office of the Senior Vice Chancellor for Health Sciences, and the UPMC Hillman Cancer Center under award P30CA047904. This study used high performance research computing core (RRID: SCR_022735) at the University of Pittsburgh Center for Research Computing supported by NIH S10OD028483 (to AVL). This research was supported in part by the University of Pittsburgh Organoid Research Core Facility (RRID: SCR_025698).

## Author Contributions

Conceptualization: AVL, SO, NC, JMA, JH

Methodology: NC, JH, JMA, SL, RL, JZ, DDB, AC, RK, DJSJM, REW III, TDN, PO, JH, HO, LR, JN, GCT, PFM, EJD, IZ, SO, AVL

Investigation: NC, JH, JMA, SL, RL, JZ, DDB, AC, RK, DJSJM, REW III, TDN, PO, JH, HO, LR, JN, DP, IZ, SO, AVL

Visualization: NC, JH, RL, SL, JZ, AC, RK, IZ, SO, AVL

Funding Acquisition: AVL, SO, NC, TDN

Project Administration: NC, AVL, SO, IZ, GCT, PFM, EJD, MTL Supervision: AVL, SO, IZ, GCT, PFM, EJD, MTL, NC

Writing – Original Draft: NC, AVL, SO Writing – Review & Editing: All authors.

## Data and Materials Availability

All raw RNA sequencing data are available on the Gene Expression Omnibus through accession numbers:

- GSE276755 (human tumor and tumor-adjacent bulk RNA-seq)
- GSE276757 (rat tumor bulk RNA-seq)
- GSE276759 (rat tumor whole exome sequencing)
- GSE276758 (rat tumor single-nuclei RNA-seq)

