## Supplementary Material for "Systemic and local chronic inflammation and hormone disposition promote a tumor-permissive environment for breast cancer in older women"

This PDF file includes:

Figs. S1 to S16
Tables S1 to S3

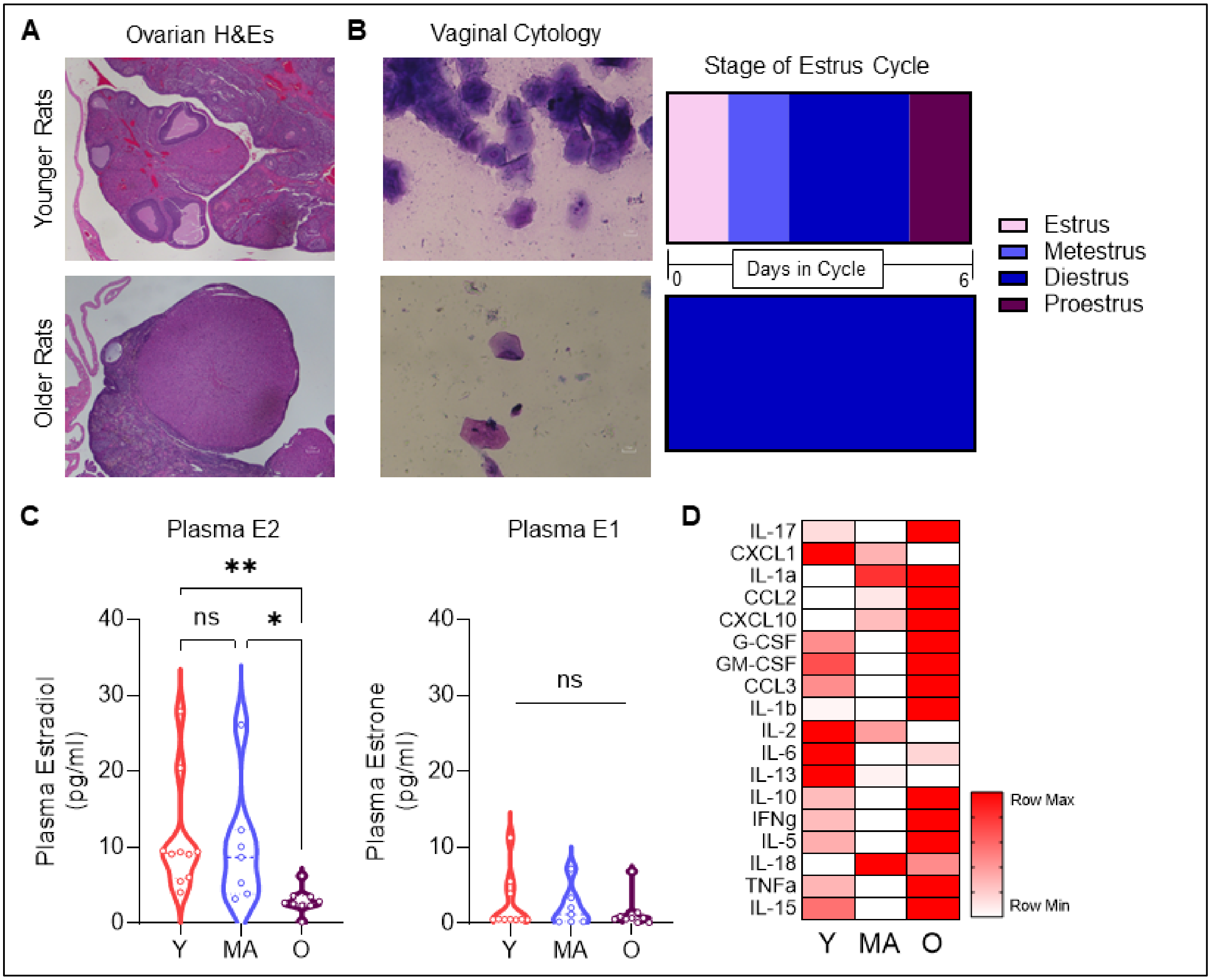


**Fig. S1: Assessment of physiological aging and estropause in F344 rats that were not exposed to chemical carcinogens.** (A) Ovarian H&E-stained slides from younger and older rats. Ovaries from younger rats (~4mo old) showed active follicle formation and cycling while the ovaries from older rats (20-22mo old) showed afollicular ovaries. (B) Representative images from vaginal cytology, in which n = 6 younger and n = 6 older rats were swabbed for 14 consecutive days. Younger rats were actively cycling every 5-6 days while the older rats were in a constant state of diestrus. (C) Analysis of circulating estradiol (E2) and estrone (E1) in young (~4mo old), middle-aged (12-13mo old), and older (20-22mo old) showed a significant decline in circulating E2 in the older rats, consistent with the human phenotype of menopause. (D) Circulating inflammation panel on the same groups of rats showed age-related chronic inflammatory phenotype in older rats.

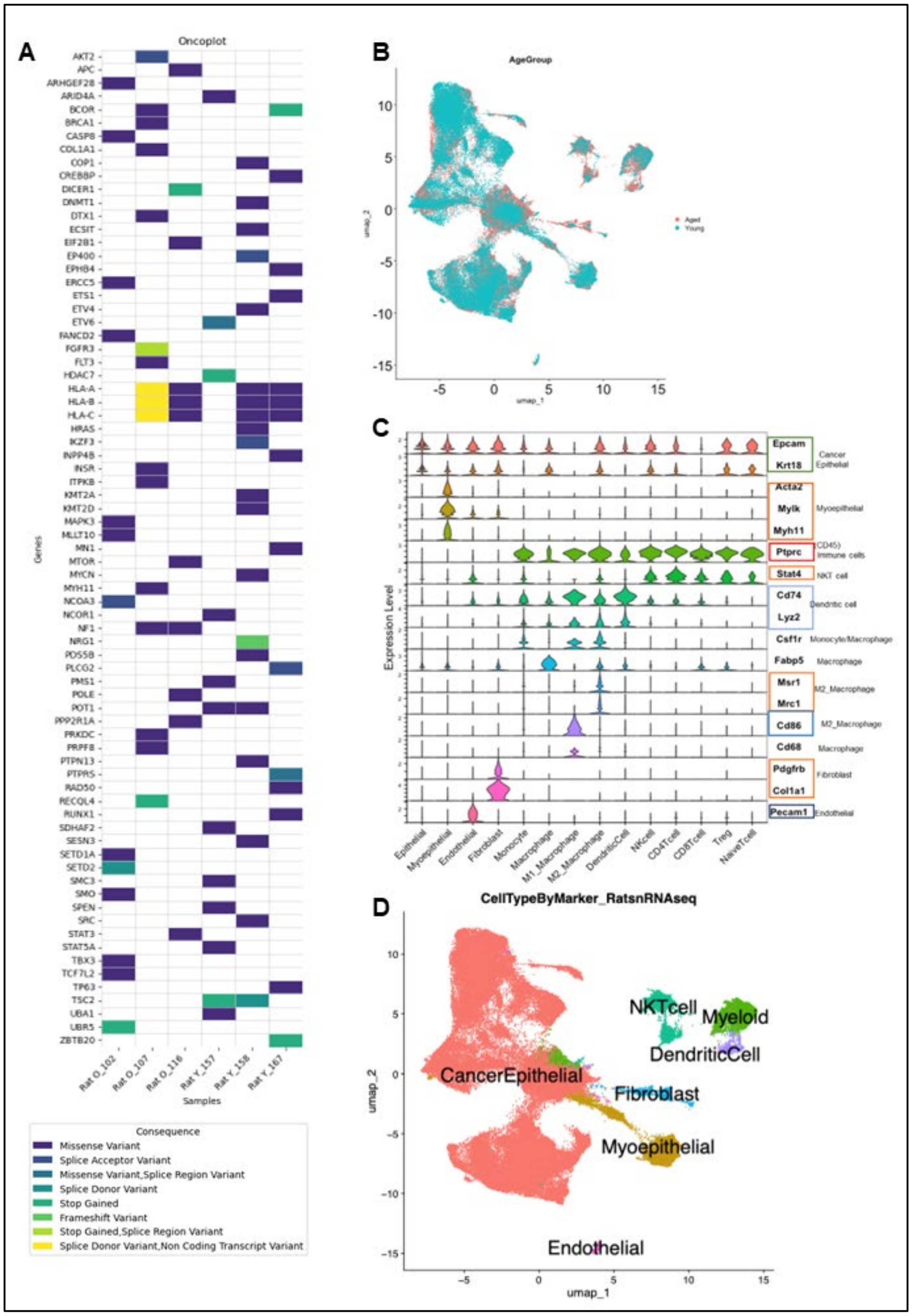


**Fig. S2: Sequencing studies of rat tumors show random mutation patterns and little genomic differences between tumors from younger and older rats.** (A) Oncoplot for the three tumors from younger rats (labeled with tumor ID and “Y”) and three tumors from older rats (labeled with tumor ID and “O”) displaying a broader group of genes. Plot shows random mutations induced from the carcinogens without a pattern related to age of the host. (B) UMAP from snRNA-seq data showing overlap between cells from tumors from young and older rats. (C) Marker genes used to identify cell types within the TME from snRNA-seq data. (D) UMAP showing the identified cell types.

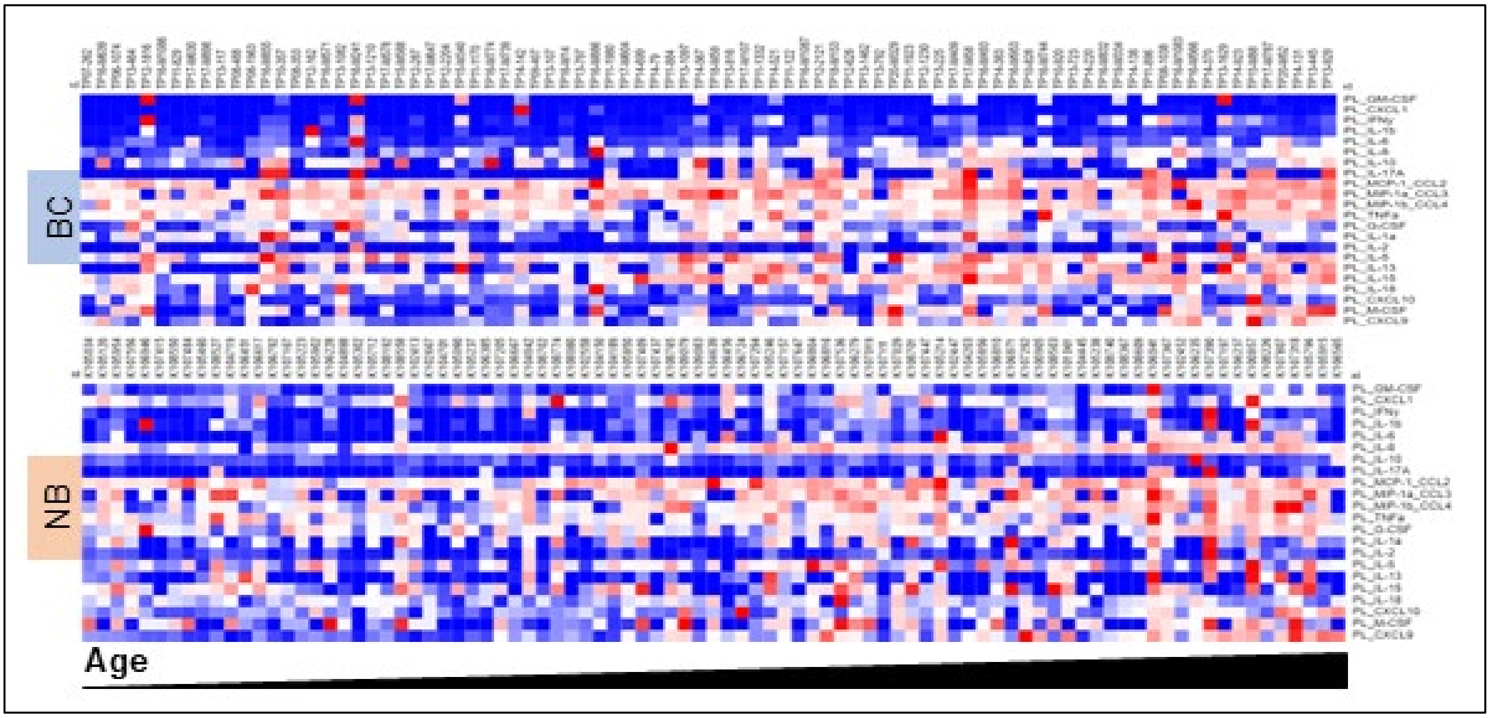


**Fig. S3: Heatmaps displaying the circulating inflammatory environment.** Heatmap shows all patient (with ER+ breast cancer, “BC”) and all donor (from the Komen Tissue Bank, “NB”) plasma analyzed using the 22-marker Luminex panel. Patients and donors are arranged according to chronological age.

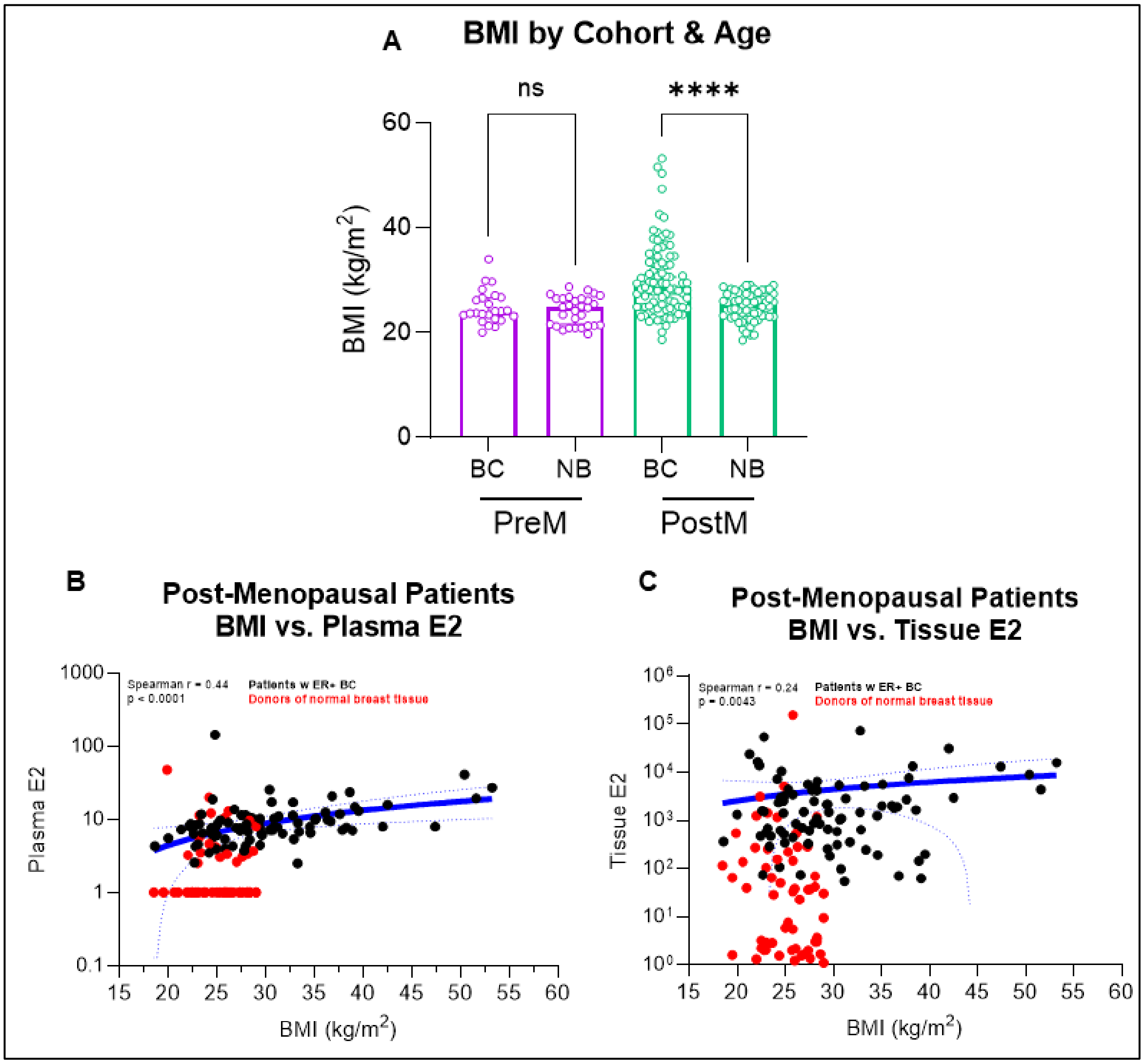


**Fig. S4: Association between host BMI estrogen levels.** (A) BMI comparison between the patients with ER+ breast cancer and the donors from the Komen Tissue Bank. **** indicates p-value < 0.0001. (B) BMI was positively associated with circulating E2 level in post-menopausal patients and donors (Spearman r = 0.44, p < 0.0001; linear regression p = 0.0113 for slope significantly nonzero). (C) BMI was weakly associated with tissue levels of E2 in post-menopausal patients and donors (Spearman r = 0.24, p = 0.0043; linear regression p = 0.37 for slope significantly nonzero).

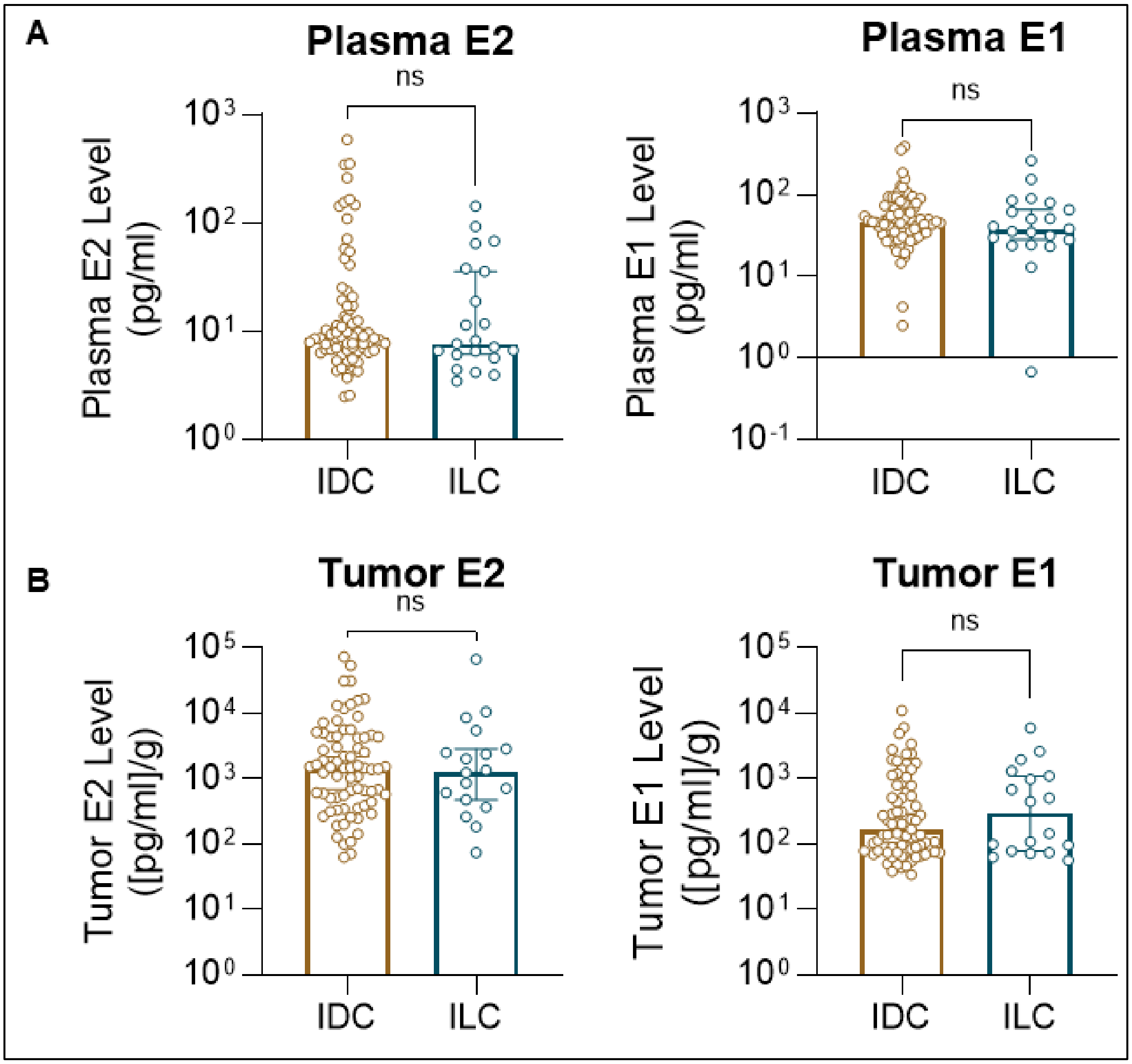


**Fig. S5: Association between estrogen levels and tumor histology.** (A and B) Plasma E1 and E2 and tumor E1 and E2 were not significantly different between patients with IDC and ILC.

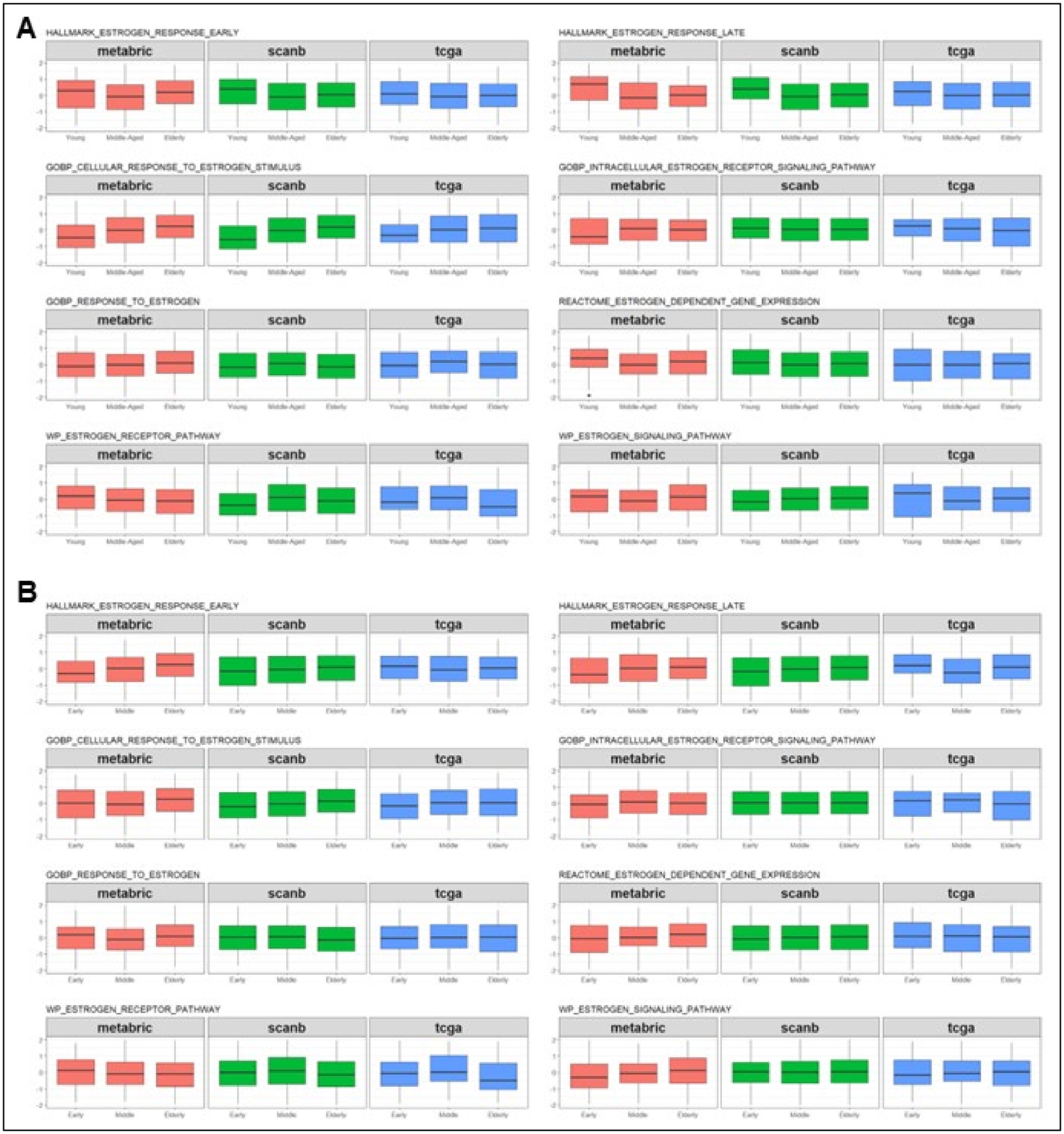


**Fig. S6: A panel of estrogen-related pathways analyzed through the mutual information concordance analysis (MICA) pipeline.**
(A) This analysis used the standard age cutoffs (Young, 35-45yo; Middle-Aged, 55-69yo; and Older, ≥ 70yo) to examine changes in estrogen-pathway activation across age groups. There were no significant changes across ages in any of the analyzed pathways. (B) This analysis only included post-menopausal patients and again found no significant differences in changes in estrogen-related pathways across ages (Early PostM, 55-60yo; Middle PostM, 61-70; Elderly, ≥ 70yo).

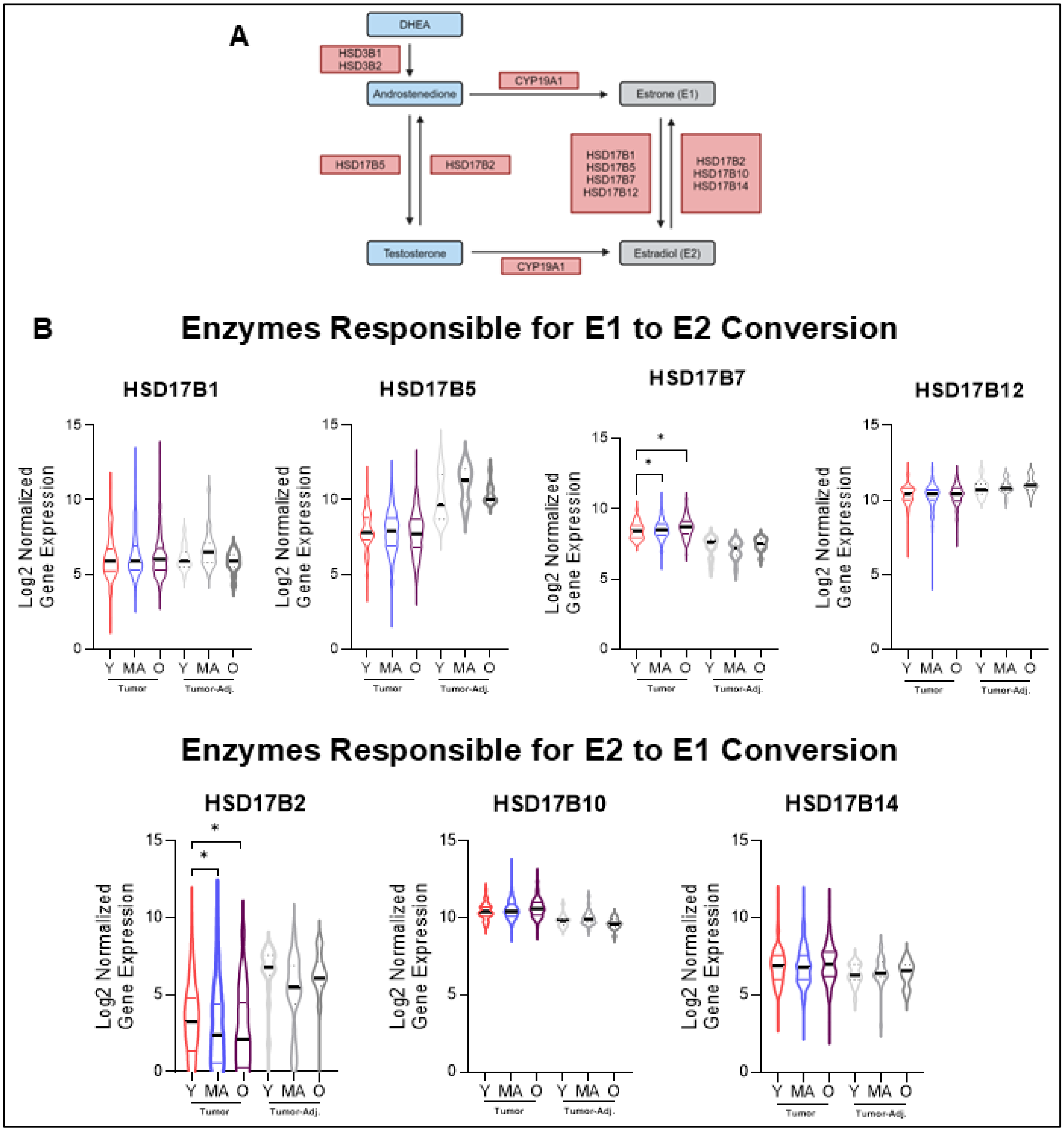


**Fig. S7: Analysis of hydroxysteroid enzymes.** (A) Schematic depicting the hydroxysteroid dehydrogenase (HSD) enzymes responsible for E1 to E2 conversion and E2 to E1 conversion. (B) Analysis of HSD enzyme gene expression levels across ages in tumor and tumor-adjacent tissue using the TCGA database.

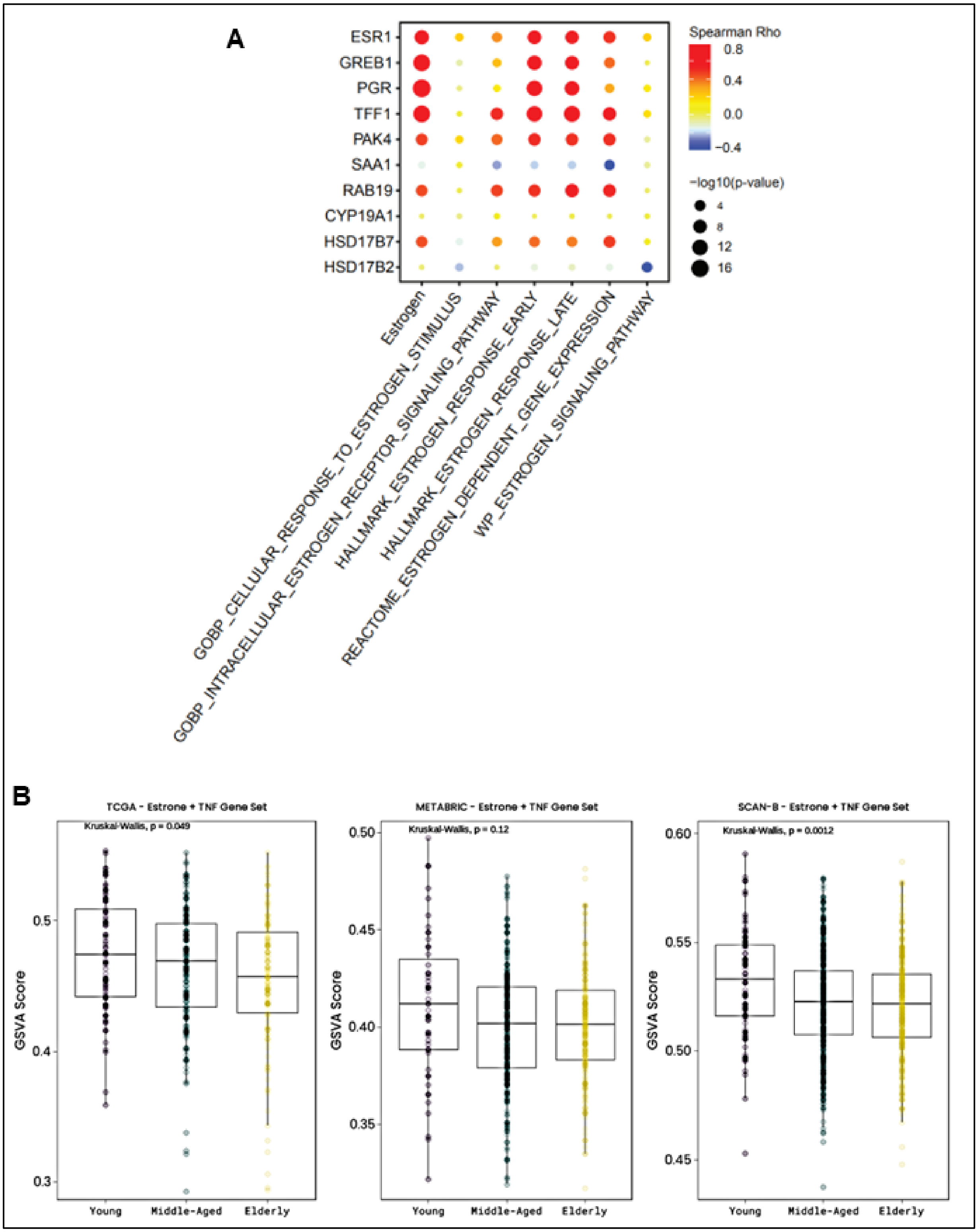


**Fig. S8: Analysis of estrogen-related pathways across ages.** (A) Correlation dot plot using the bulk RNA-seq data generated in this study showing correlation between several estrogen-related pathways vs. estrogen-related genes and enzymes. (B) Analysis of an Estrone+TNF signature (ref. 28) across ages in TCGA, METABRIC, and SCAN-B.

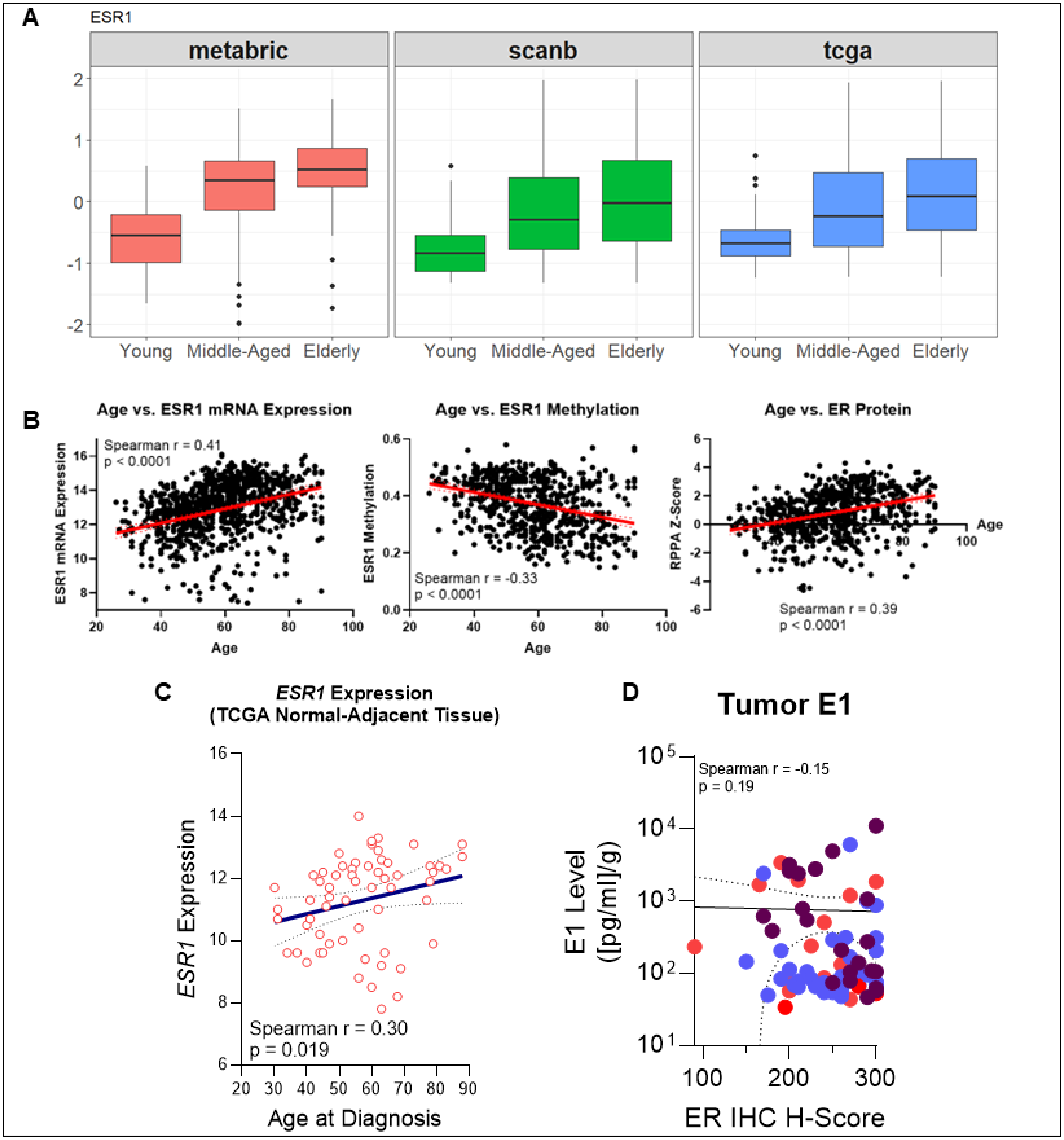


**Fig. S9: ESR1 changes with age.** (A) ESR1 expression levels across age in the TCGA, METABRIC, and SCAN-B datasets. (B) Scatter plots showing Age vs. ESR1 mRNA expression, ESR1 methylation, and ER protein (all using the TCGA database). (C) Scatter plot showing Age vs. ESR1 expression in TCGA normal tumor-adjacent tissues. (D) Scatter plot showing ER IHC H-score vs. tumor E1 levels.

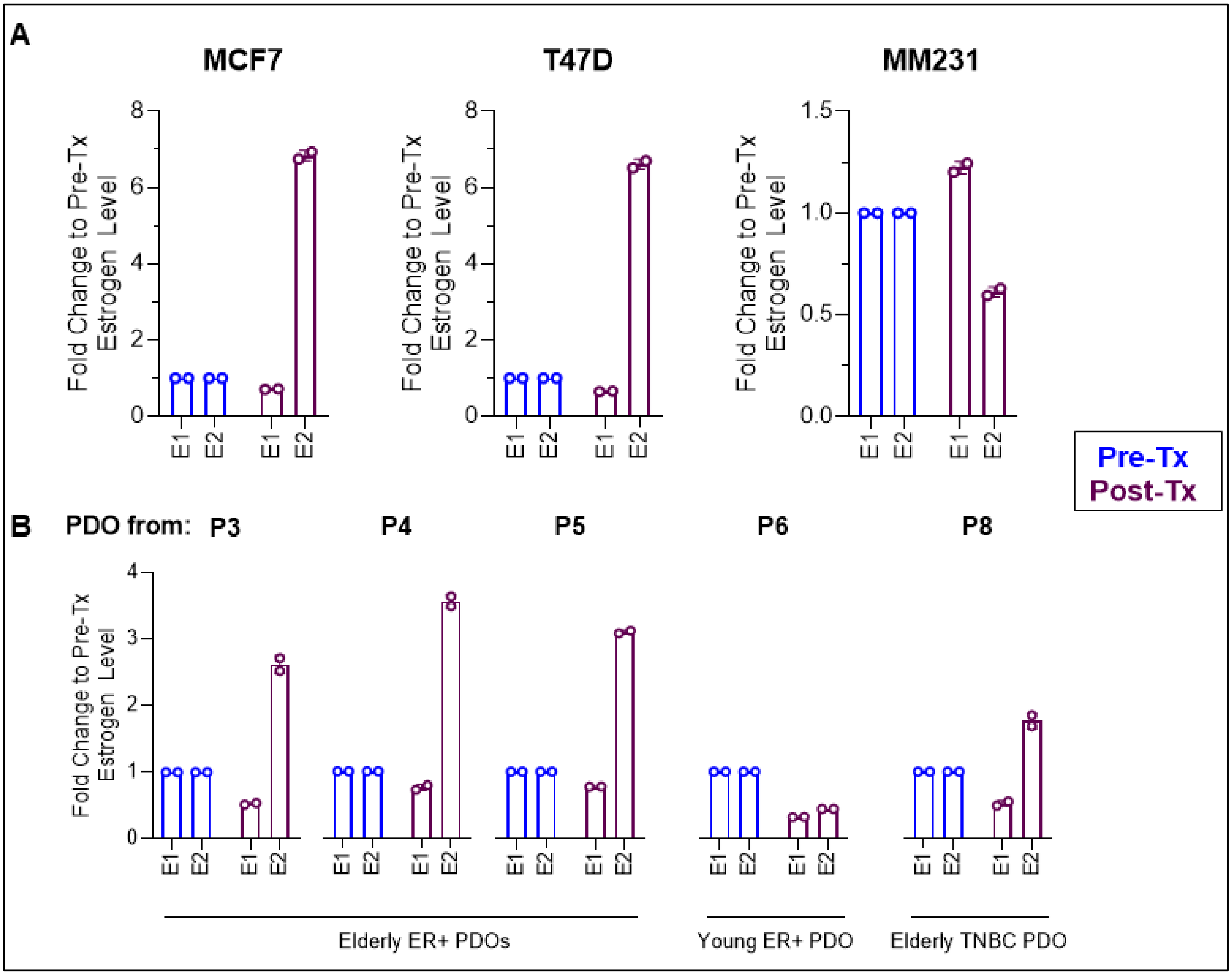


**Fig. S10: E1 treatment of cell lines and PDOs.** (A and B) We treated cell lines (MCF7, T47D, and MM231) as well as a panle of patient-derived organoids (PDOs) with E1 and evaluated the degree to which that E1 was converted to E2. All post-treatment data is normalized to the pre-treatment levels. PDOs from Patient 3 (P3), 4 (P4), 5 (P5), 6 (P6), and 8 (P8) were used in this experiment (see Table S3 in this document for full clinical characteristics of the PDOs used in this study). P3, P4, and P5 were PDOs derived from older patients with ER+ breast cancer; P6 was a PDO derived from a younger, pre-menopausal patient with ER+ breast cancer; and P8 was a PDO derived from an older patient with TNBC.

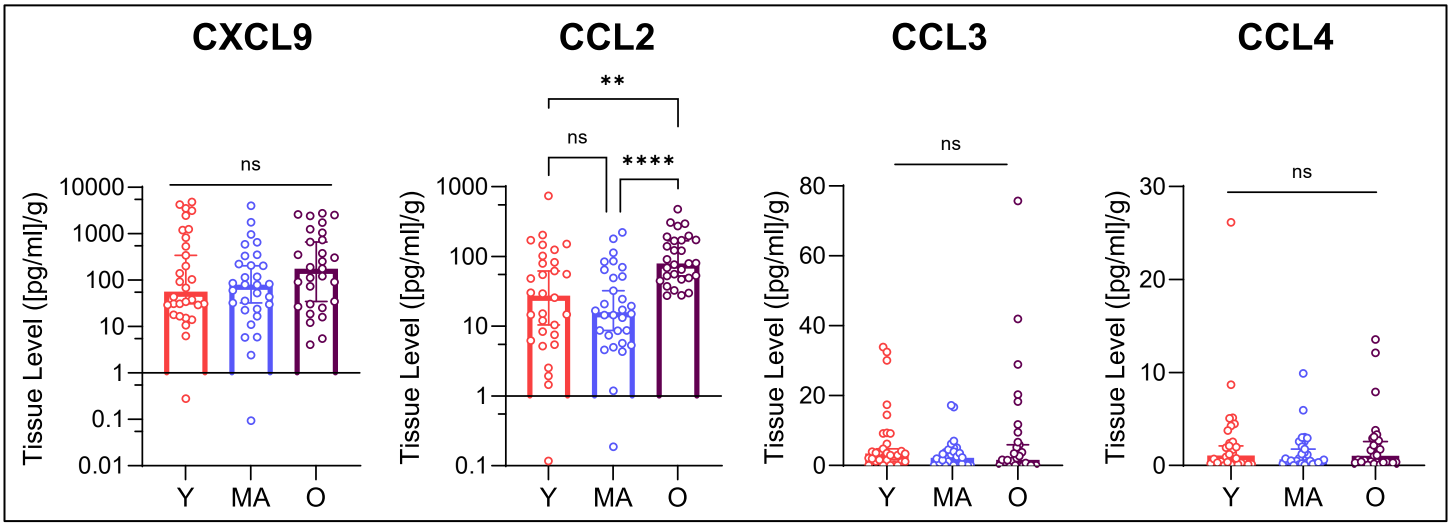


**Fig. S11: Chemokine levels across age from donors of normal breast tissue.** Data shows measured CXCL9, CCL2, CCL3, and CCL4 across age as measured from the tissue lysates from normal breast tissue from donors of the Komen Tissue Bank.

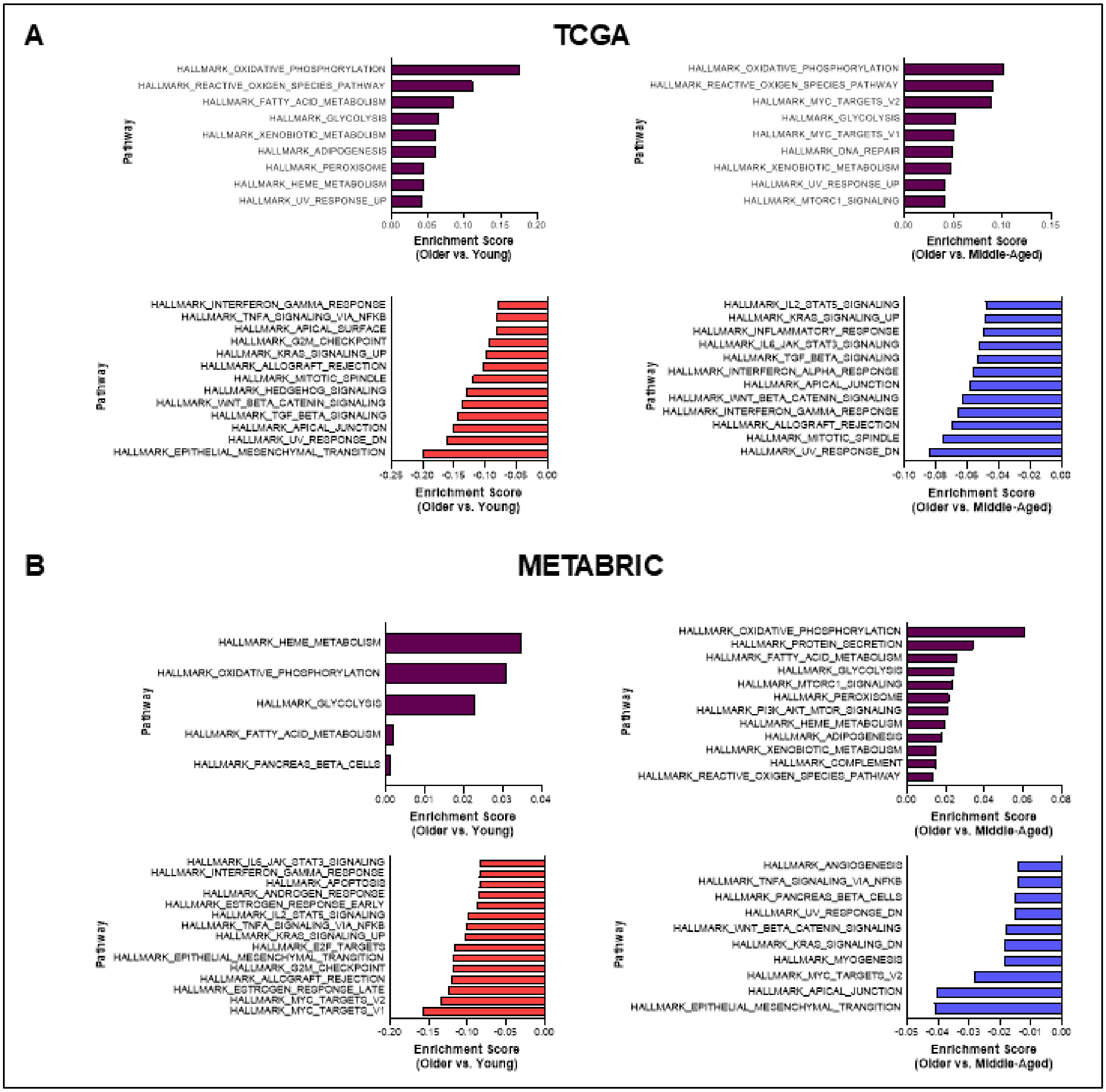


**Fig. S12: Pathway enrichment comparisons using TCGA and METABRIC data.** (A) Shows top and bottom enriched pathways in older patients compared to both middle-aged and younger patients in TCGA. (A) Shows top and bottom enriched pathways in older patients compared to both middle-aged and younger patients in METABRIC.

A

| Age_Group | Case ID | Age | Grade | Histological Subtype | Stage | ER | PR | HER2 IHC | HER2 ISH (ratio) | Ki67 | Treatment status | Notable Pathological features |
|---|---|---|---|---|---|---|---|---|---|---|---|---|
| O | 4290 | 88 | 2 | IDC | pT4b, Nx | 90% 3+ | 30% 2+ | 1+ | Non-Amplified | 10% | Naïve | Locally advanced, skin and chest wall muscle involvement. |
| O | 4067 | 85 | 2 | IDC | pT2, N1(sn), Mx | 100% 3+ | 95% 3+ | 1+ | Non-Amplified | 3-4% | Naïve | Associated low grade DCIS and focal perineural invasion. |
| O | 3948 | 82 | 3 | IDC | pT2, N2a | 90% 2-3+ | 80% 2+ | 0 | Non-Amplified | 10% | Naïve | Associated LCIS, with LVI and perineural invasion |
| MA | 4463 | 58 | 2 | IDC | pT3, N1, Mx | 100% 2-3+ | 80% 2-3+ | 0 | Non-Amplified | 50% | Naïve | IDC with areas of lobular-like growth pattern, but is E-cadherin positive. Associated low through high grade DCIS and LVI. |
| MA | 4040 | 57 | 3 | IDC | pT2, N0 | 95% 3+ | 95% 2-3+ | 0 | Non-Amplified | > 50% | Naïve | Associated high grade DCIS. |
| MA | 4471 | 55 | 2 | ILC | pT3, pN0 (i+) | 100% 3+ | 100% 3+ | 0 | Non-Amplified | 20% | Naïve | None. |
| MA | 4461 | 54 | 2 | IDC | pT3, N1a, Mx | 95% 3+ | ~5% 3+ | 2+ | Non-Amplified | 15% | Naïve | Associated intermediate to high grade DCIS, LVI and perineural invasion. |
| Y | 3941 | 50 | 2 | IDC | pT1c, N1a, Mx | 90% 3+ | 90% 3+ | 2+ | Non-Amplified | 10% | Naïve | Multifocal tumour with associated high grade DCIS |
| Y | 4535 | 47 | 2 | ILC | pT2, pN0 (i+),Stage IIB | 95% 3+ | 70% 2+ | 2+ | Non-Amplified | 10% | Naïve | None. |
| Y | 4530 | 42 | 2 | IDC | pT3, pN3, pMx, Stage IIIA | 95% 2+ | 95% 3+ | 1+ | Non-Amplified | 5% | Naïve | Multifocal tumour with associated high grade DCIS and LVI. |

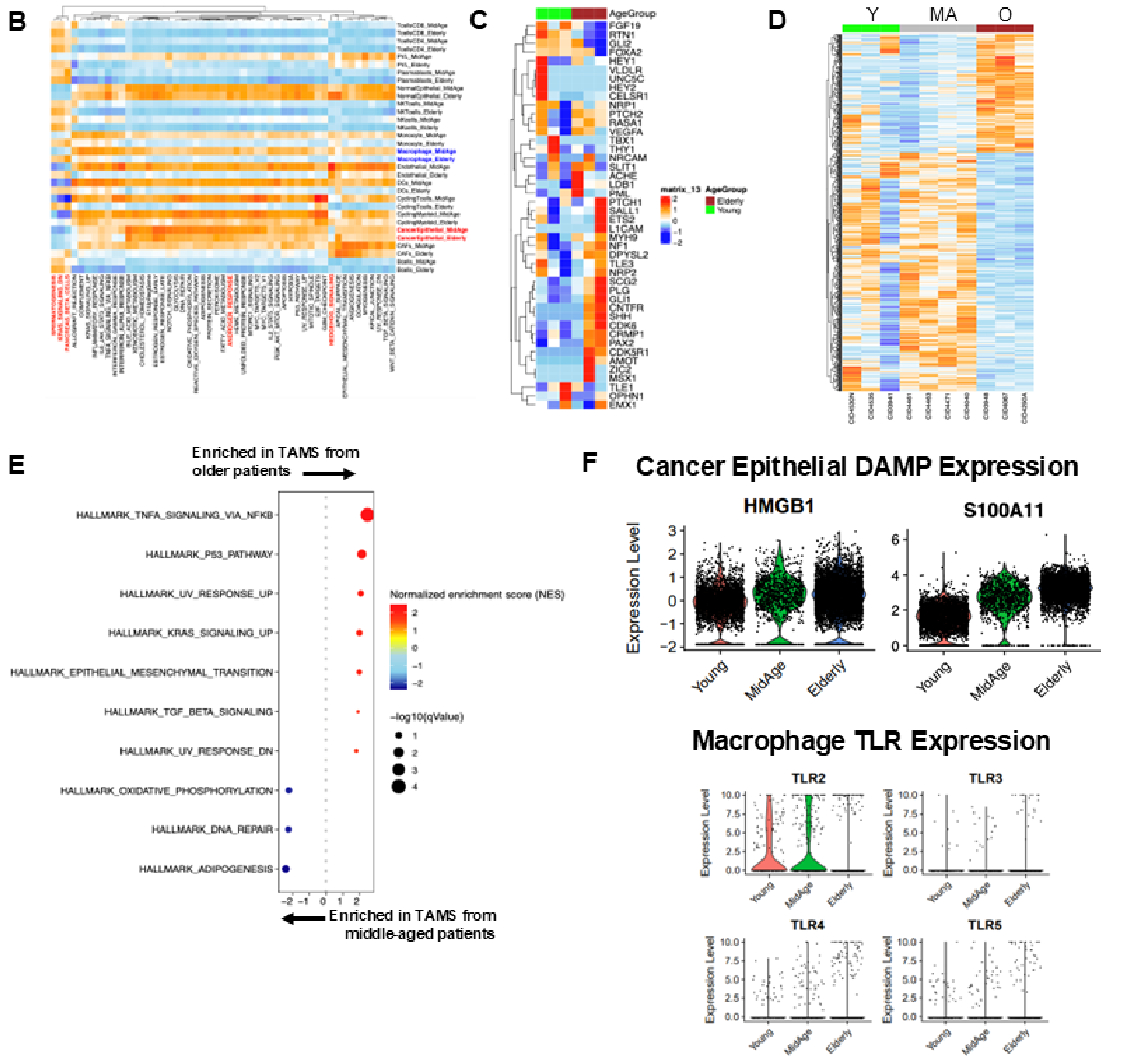


**Fig. S13: Analysis of macrophages in older patients using publicly-available scRNA-seq data.** (A) We analyzed data derived from patients in ref. 68, only including those with treatment-naïve, ER+/HER2- tumors. Table shows the patients that were included. (B) Heatmap showing transcriptomic differences in immune cells from older and middle-aged patients. (C) Heatmap showing expression of Hedgehog pathway genes in macrophages from the tumors of older and younger patients. (D) Heatmap showing DEGs between macrophages in younger, middle-aged, and older tumors. (E) Pathway analysis of the DEGs in macrophages from older and middle-aged patients. (F) Expression of DAMP molecules from cancer epithelial cells across age groups and macrophage toll-like receptor (TLR) expression across age groups.

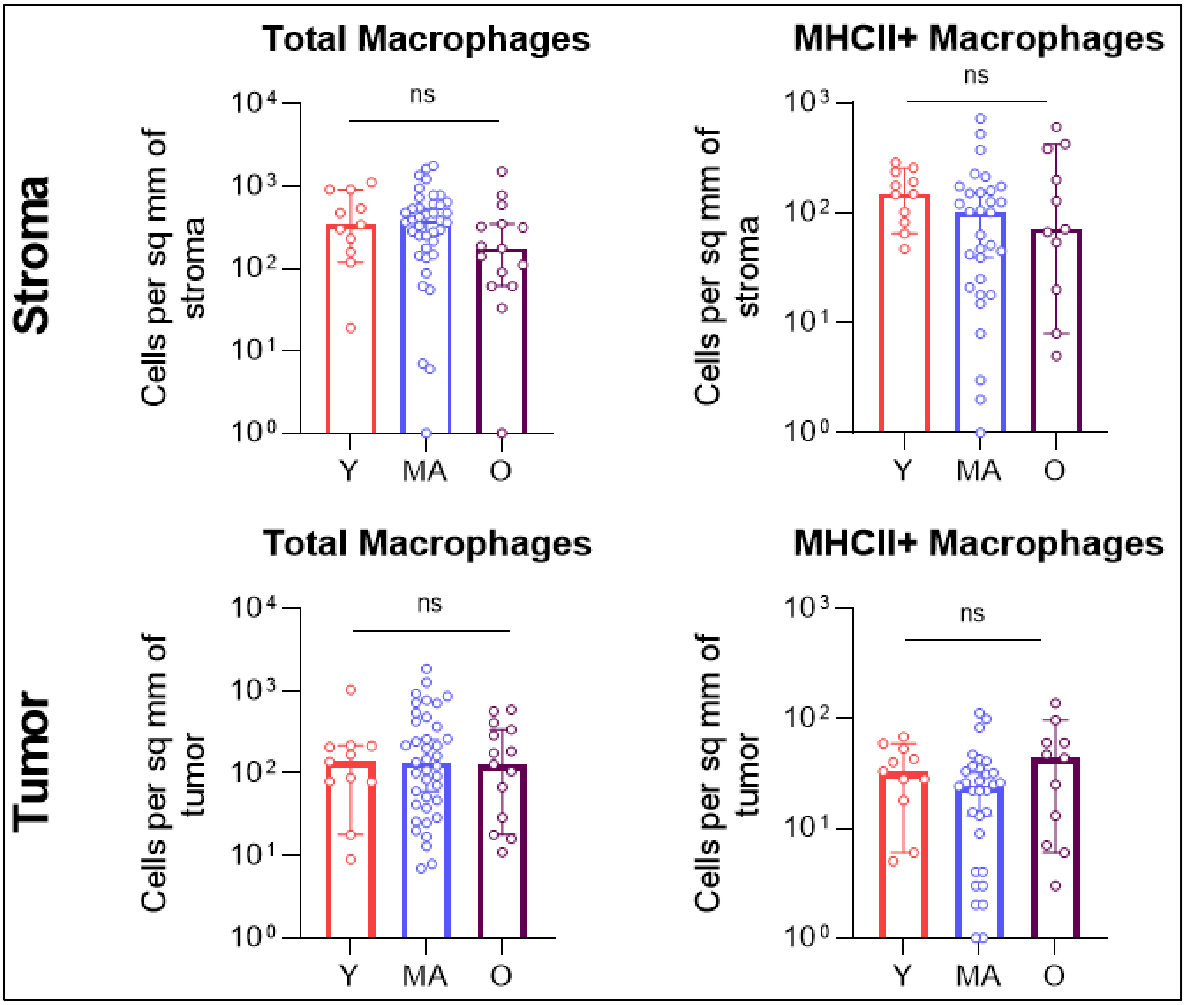


**Fig. S14: mIHC data showing total macrophage and MHCII+ macrophage quantification.** Using mIHC data from ref. 35 (Onkar et al), we reanalyzed the data according to age group, showing little difference in total macrophage and MHCII+ macrophage levels across age groups.

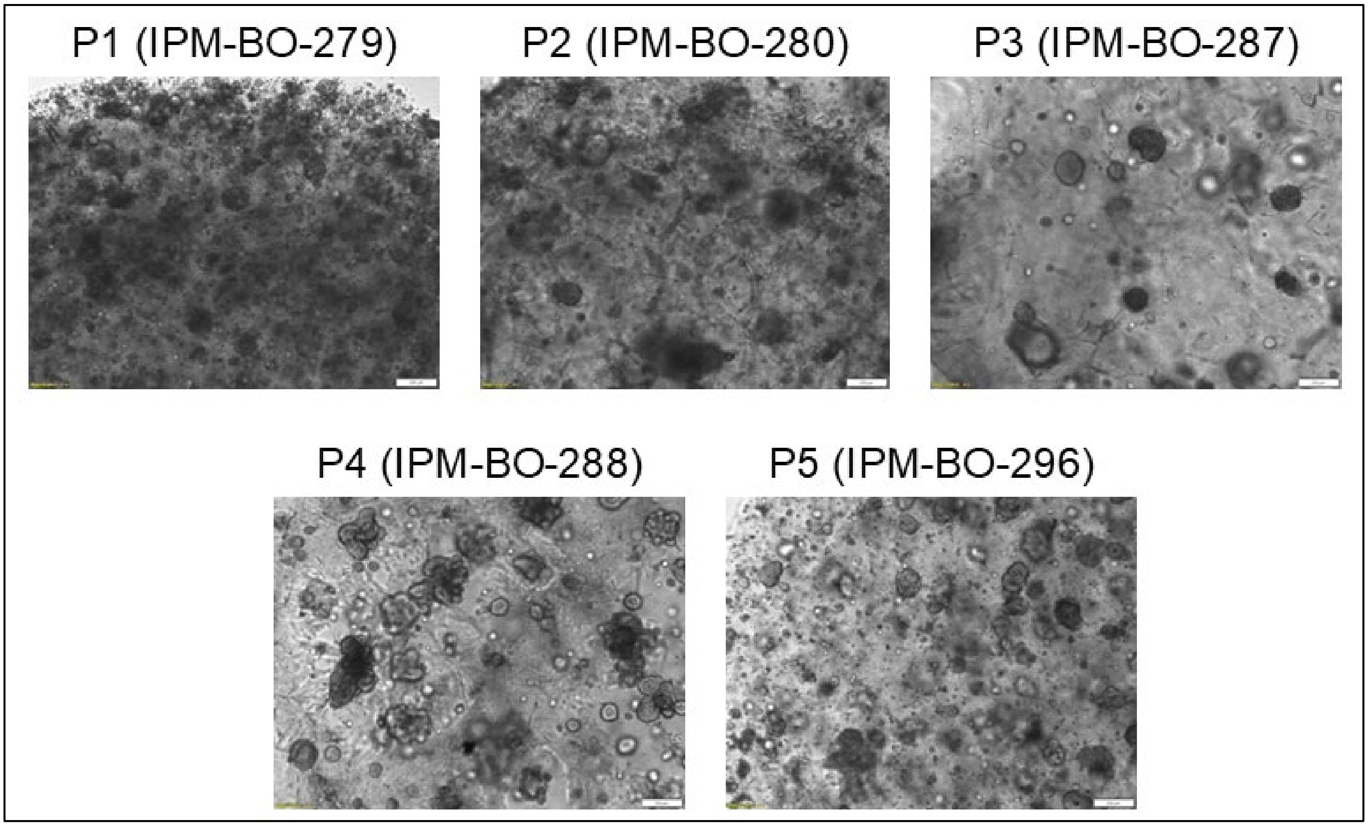


**Fig. S15: Brightfield images of just-in-culture patient-derived organoids (PDOs).** PDOs freshly in culture a diverse milieu of cells (including tumor cells, endothelial cells, lymphocytes, and fibroblasts. Images take at 4x magnification; scale bar represents 200µm.

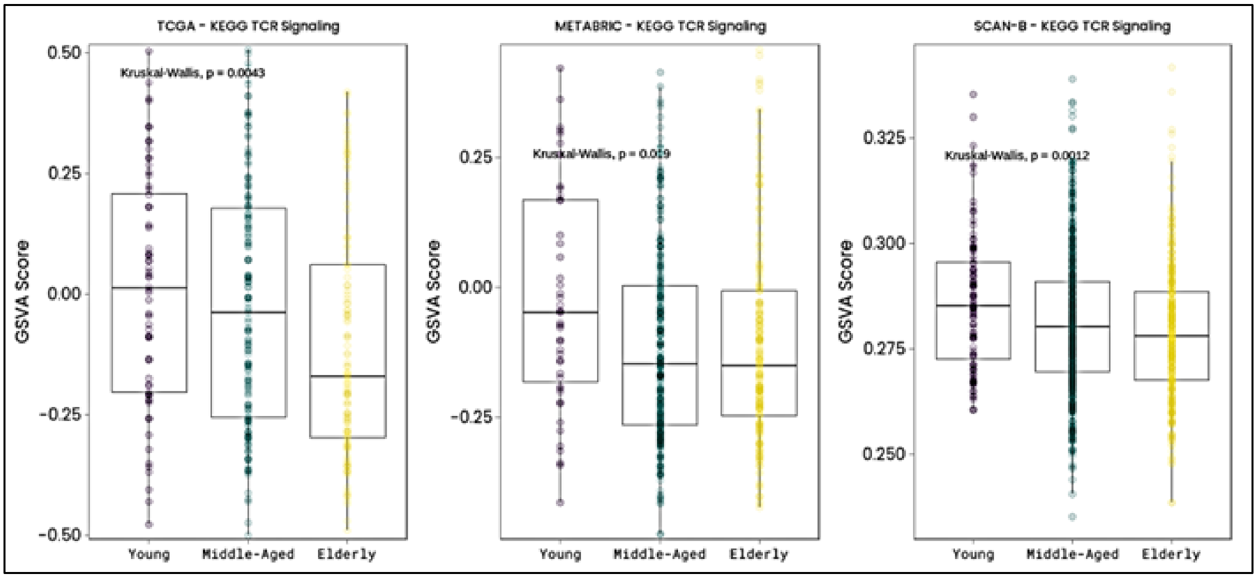


**Fig. S16: KEGG TCR signaling pathway enrichment across age.** Plots show the pathway enrichment levels of the KEGG TCR signaling pathway across the three age groups, with the lowest pathway enrichment level in older patients.

| | Young Patients N = 25 | Middle-Aged Patients N = 57 | Older Patients N = 33 | P-value |
|---|---|---|---|---|
| **Median age at diagnosis, years (IQR)** | 39 (37-42) | 62 (58.5-65.5) | 76 (72.5-79) | < 0.0001 |
| **Median tumor size, mm (IQR)** | 21 (17-25) | 21 (15-37.5) | 21 (12-29) | 0.36 |
| **Median BMI, kg/m$^2$ (IQR)** | 23.9 (22.7-26.7) | 29.4 (25.1-34.3) | 27.5 (23.3-33.9) | 0.0001 |
| **Grade at Diagnosis, n (%)** | | | | 0.38 |
| I | 3 (12.0%) | 3 (5.3%) | 2 (6.1%) | |
| II | 11 (44.0%) | 36 (63.1%) | 23 (69.7%) | |
| III | 10 (40.0%) | 15 (26.3%) | 8 (24.2%) | |
| Unknown | 1 (4.0%) | 3 (5.3%) | 0 (0%) | |
| **Stage at Diagnosis, n (%)** | | | | 0.66 |
| I | 12 (48.0%) | 24 (42.1%) | 10 (30.2%) | |
| II | 6 (24.0%) | 17 (29.8%) | 13 (39.4%) | |
| III | 5 (20.0%) | 11 (19.3%) | 6 (18.2%) | |
| Unknown | 2 (8.0%) | 5 (8.8%) | 4 (12.1%) | |
| **Histological Subtype, n (%)** | | | | 0.67 |
| IDC / NST | 16 (64.0%) | 33 (57.9%) | 23 (69.7%) | |
| ILC | 5 (20.0%) | 10 (17.5%) | 6 (18.2%) | |
| Other / Multiple Types Present | 4 (16.0%) | 14 (24.6%) | 4 (12.1%) | |
| **Menopausal Status, n (%)** | | | | < 0.0001 |
| Pre-Menopausal | 23 (92.0%) | 4 (7.0%) | 0 (0%) | |
| Post-Menopausal | 2 (8.0%) | 53 (93.0%) | 33 (100%) | |
| **Any History of Smoking, n (%)** | | | | 0.90 |
| Any prior / current use | 10 (40.0%) | 27 (47.3%) | 15 (45.5%) | |
| Never Used | 14 (56.0%) | 30 (52.7%) | 18 (54.5%) | |
| Unknown | 1 (4.0%) | 0 (0%) | 0 (0%) | |
| **Any History of EtOH Use, n (%)** | | | | 0.51 |
| Any prior / current use | 3 (12.0%) | 8 (14.0%) | 2 (6.1%) | |
| No significant history | 21 (84.0%) | 49 (86.0%) | 31 (93.9%) | |
| Unknown | 1 (4.0%) | 0 (0%) | 0 (0%) | |
| **Patient Race** | | | | 0.47 |
| White | 23 (92.0%) | 52 (91.2%) | 31 (94.0%) | |
| Black | 2 (8.0%) | 5 (8.8%) | 1 (3.0%) | |
| Other | 0 (0%) | 0 (0%) | 1 (3.0%) | |
| **Patient Ethnicity** | | | | 0.08 |
| Non-Hispanic | 22 (88.0%) | 55 (96.5%) | 33 (100%) | |
| Hispanic | 3 (12.0%) | 2 (3.5%) | 0 (0%) | |

**Table S1:** Patient and tumor characteristics from the 115 patients with ER+ breast cancer whose specimens (blood, tumor tissue, and tumor-adjacent tissue) were used in this study.

| | | Young Donors N = 30 | Middle-Aged Donors N = 30 | Older Donors N = 29 | P-value |
|---|---|---|---|---|---|
| **Median age at time of tissue donation, years (IQR)** | | 37 (36-38.5) | 58 (58-59) | 74 (71.5-77) | < 0.0001 |
| **Median BMI, kg/m$^2$ (IQR)** | | 24.7 (21.6-26.53) | 24.4 (22.5-26.8) | 25.8 (23.8-28.3) | 0.09 |
| **Age at Menarche, years (IQR)** | | 13 (11.8-14) | 13 (12.8-14) | 13 (12-14) | 0.25 |
| **Breast Cancer Risk Assessment Gail Score, median (IQR)** | | 11.3 (9.7-15.7) | 8.9 (7.5-13.6) | 4.9 (3.9-8.4) * | < 0.0001 |
| **Number of Prior Pregnancies** | | | | | 0.11 |
| | 0 | 8 (26.7.0%) | 5 (16.7%) | 1 (3.5%) | |
| | 1 | 5 (16.7%) | 3 (10.0%) | 3 (10.3%) | |
| | ≥ 2 | 17 (56.6%) | 22 (73.3%) | 25 (86.2%) | |
| **Any Prior HRT Use [#]** | | | | | < 0.0001 |
| | Yes | 0 (0%) | 8 (26.7%) | 15 (51.7%) | |
| | No | 29 (96.7%) | 22 (73.3%) | 14 (48.3%) | |
| | Unknown | 1 (3.3%) | 0 (0%) | 0 (0%) | |
| **Patient Race** | | | | | 0.18 |
| | White | 28 (93.3%) | 30 (100%) | 29 (100%) | |
| | Black | 2 (6.7%) | 0 (0%) | 0 (0%) | |
| **Patient Ethnicity** | | | | | 0.35 |
| | Non-Hispanic | 26 (86.7%) | 26 (86.7%) | 28 (96.6%) | |
| | Hispanic | 4 (13.3%) | 4 (13.3%) | 1 (3.4%) | |

* Indicates five missing values for Gail Score within the older donor group.
[#] No donors with prior HRT use were taking HRT at the time of tissue donation. Information is not available as to the time off of HRT prior to tissue/blood donation.

**Table S2:** Characteristics from the 89 donors (from the Komen Tissue Bank at Indiana University) whose specimens (blood and non-tumor, normal breast tissue) were used in this study. * Indicates five missing values for Gail Score within the older donor group.

| Label | Patient Age | Tumor Receptors | Ki67 | Histology | NS | Pathologic Stage | UPMC Institute of Precision Medicine PDO ID |
|---|---|---|---|---|---|---|---|
| Patient 1 (P1) | 88 | ER 100, PR 12, HER2 0 | 90% | IDC | 9/9 | pT3N1 | IPM-BO-279 |
| Patient 2 (P2) | 81 | ER 300, PR 1, HER2 0 | 30% | IDC | 6/9 | pT3N0 | IPM-BO-280 |
| Patient 3 (P3) | 78 | ER 290, PR 150, HER2 1+ | 55% | IDC | 7/9 | pT2N1 | IPM-BO-287 |
| Patient 4 (P4) | 76 | ER 300, PR 300, HER2 0 | 10% | mDLC | 6/9 | pT1cN0 | IPM-BO-288 |
| Patient 5 (P5) | 83 | ER 250, PR 300, HER2 2+ NA | 65% | IDC | 6/9 | pT2N0 | IPM-BO-296 |
| Patient 6 (P6) | 43 (PreM) | ER 290, PR 290, HER2 0 | 10% | IDC | 6/9 | pT1cN1 | IPM-BO-056 |
| Patient 7 (P7) | 50 (PreM) | ER 290, PR 290, HER2 0 | 5% | IDC | 6/9 | pT2N2 | IPM-BO-126 |
| Patient 8 (P8) | 79 | ER 0, PR 0, HER2 0 | 65% | IDC | 6/9 | pT3N0 | IPM-BO-229 |

**Table S3:** Age and tumor characteristics for the patient derived organoids (PDOs) that were used in this study. All PDOs were collected from specimens at the time of surgery; all patients did not receive neoadjuvant systemic therapy prior to PDO generation. We note that while P8 would have been eligible for neoadjuvant therapy with the Keynote-522 regimen, this patient had a number of comorbidities and opted for upfront surgery.
